# Decoding natural astrocyte rhythms: dynamic actin waves result from environmental sensing by primary rodent astrocytes

**DOI:** 10.1101/2021.09.13.460152

**Authors:** Kate M. O’Neill, Emanuela Saracino, Barbara Barile, Nicholas J. Mennona, Maria Grazia Mola, Spandan Pathak, Tamara Posati, Roberto Zamboni, Grazia P. Nicchia, Valentina Benfenati, Wolfgang Losert

## Abstract

Astrocytes are key regulators of brain homeostasis, an important physiological process that includes but is not limited to buffering of extracellular K^+^, equilibrating osmotic gradients, regulating pH, uptake of neurotransmitters, and releasing growth factors – all of which are essential for proper cognitive function. While previous studies have revealed how specific molecular components of the astrocytic cytoskeleton affect the efficacy of these homeostatic processes, none have studied how homeostasis is linked to the excitable systems character of the cytoskeleton. As recently discovered, excitability of the actin cytoskeleton manifests in second-scale dynamic fluctuations and acts as a sensor of chemo-physical environmental cues. Here we find that homeostatic regulation may be more active than previously thought, involving the excitable dynamics of actin in certain subcellular regions, especially near the cell boundary. Our results further indicate that actin dynamics concentrates into “hotspot” regions that selectively respond to certain chemo-physical stimuli, specifically the homeostatic challenges of ion or water concentration increases. Substrate topography makes the actin dynamics of astrocytes weaker. Superresolution images demonstrate that surface topography is also associated with predominant perpendicular alignment of actin filaments near the cell boundary whereas flat substrates result in an actin cortex mainly parallel to the cell boundary. Additionally, co-culture with neurons increases both the probability of actin dynamics and the strength of hotspots. The excitable systems character of actin thus makes astrocytes direct participants in neural cell network dynamics.

## Introduction

The ability of the brain to receive and compute information depends on the unique abilities of its component brain cells. After each neuronal action potential, the concentrations of ions and water are altered in the perineural and perivascular spaces. Specifically, the concentration of potassium in these areas may be as high as 60-100 mM [1–3] in pathophysiological situations. One of astrocytes’ homeostatic functions is to balance extracellular potassium concentration through potassium uptake and spatial buffering [4–12]. This process relies on the sodium/potassium pump (Na^+^/K^+^ ATPase), potassium channels from the inward rectifier superfamily, and gap junctional coupling. Moreover, this transmembrane flux of K^+^ is also accompanied by osmotically-driven water movement and intracellular calcium signaling [13–16].

With a stellate morphology [17–19], astrocytic processes contact neurons and blood vessels and therefore play critical roles in coupling the neuronal with the vascular activities of the brain [20]. Perisynaptic astrocytic processes (PAPs) and perivascular astrocytic processes (PvAPs) represent functional microdomains and are enriched with water and ion channels that allow astrocytes to precisely control the extracellular environment through continuous sensing and balancing of local ionic changes (between neurons for PAPs and at the blood vessel interface for PvAPs) in response to neuronal network activity [20–26]. At the same time, the gap junctional coupling between astrocytes serves as a signaling mechanism enabling synchronicity and lateral communication between astrocytes within the larger astroglia syncytia [27,28].

Though astrocytic communication via gap junctional coupling enables dynamic exchange between cells at the ionic and molecular scales [1,27–29], questions remain how these small scale dynamics couple with, modulate, and impact responses at larger length scales, such as within astroglial cell networks or the whole brain. In particular, gap junctions participate in potassium spatial buffering by aiding in the dispersion of excess K^+^ through the astroglial syncytium [1,4–7] and are also implicated in the propagation of astrocytic calcium waves, a form of extraneuronal signaling [30–34]. The efficiency of these homeostatic processes [35,36] is considered the basis for astrocytes’ involvement in cognitive functions while their impairment may cause pathologies ranging from edema to epilepsy [16].

More recently, the importance of astrocytes’ structural and functional properties has been highlighted for organ- and system-level functions, including synaptic plasticity, memory formation, learning, and regulation of sleep and metabolic activity [37,38]. Here we investigate the actin cytoskeleton as an integrator from the molecular to the systems scale due to its ability to self-assemble into dynamic networks. Indeed, experiments on fixed cells have revealed the key molecular players regulating changes to astrocyte morphology, including but not limited to Rac1 and RhoA GTPases [39–44] and the Arp2/3 complex [45], all of which play a role in promoting, maintaining, or reversing the stellate morphology typical of differentiated astrocytes [46]. Other recent studies on fixed cells have also indicated that the astrocytic actin cytoskeleton plays a role in several homeostatic processes, such as volume regulation and potassium spatial buffering [47–49].

However, much less is known about the second-scale dynamics of the actin cytoskeleton in these cellular processes. For migrating cells – such as neutrophils, metastatic cancer cells, and even microglia – actin dynamics had been mostly considered as a facilitator of changes in cell morphology. For example, a recent study in T cells demonstrated the importance of global (whole cell) actin dynamics in facilitating immunological synapse formation [50]. New lines of research, though, are also investigating the importance of actin dynamics in crucial cell processes. Dynamic actin has been shown to be important for biological function [51,52], such as surveillance of the brain parenchyma by microglia [53]. Notably, actin cytoskeletal dynamics are necessary for relaying biomechanical forces at the synapse [54], indicating that mechanical signals (in addition to the well-studied chemical and electrical signals) are critical for synaptic transmission and, therefore, cognitive function.

Moreover, recent studies across multiple cell types have demonstrated that the actin cytoskeleton is not only dynamic when a cell changes shape and moves but is also an excitable system with its own intrinsic dynamics, including sustained waves and oscillations that persist independently of changes to cell shape [55–64]. To this end, we have demonstrated that actin waves serve as sensors of electric field strength and direction in the *D. discoideum* model system [64], and we both suspect and have demonstrated that this role extends to other cell types [63]. The source of actin dynamics is thought to be twofold: i) a cytoskeletal excitable network of proteins (CEN) that drives cycles of polymerization and depolymerization typically lasting a few seconds, and ii) a signal transduction excitable network (STEN) that can self-organize these local oscillations into larger scale waves that drive cell migration [51,58,59,65]. We propose that, in astrocytes, CEN dynamics dominate, thus leading to local actin dynamics that is stationary in space and occurs on a second timescale. Therefore, CEN could carry information about homeostatic challenges that occur at the nanometer, ionic, and molecular scales.

Here we study how the actin cytoskeleton serves as the bridge from these smaller scales (e.g., ions) to larger scales (e.g., cells) and facilitates the global (cellular level) homeostatic response of astrocytes. We analyze actin dynamics using optical flow [60,66] and perform shape analysis [57,67,68] on timelapse images of live primary astrocytes transduced with actin-GFP. We demonstrate that actin dynamics occurs in astrocytes and that it can be triggered by extracellular environment modifications, such as an increase in ion concentrations or a change in culturing environment. The spatial distribution of the actin dynamics is localized near the boundary of the cell and is concentrated into active clusters we term “hotspots”.

Notably, the strength of actin dynamics changes depending on whether the exposure is to an increase in extracellular potassium or to hypotonic challenge, abbreviated as “high K^+^” and “hypotonic”, respectively. We refer to the application of these solutions to cultures as “triggering” conditions because they have been shown to mimic pathophysiological changes in the extracellular milieu similar to epilepsy [69] or edema [70] and to cause responses in astrocytes that range from swelling to calcium waves [3,48,71–78]. In addition, activity identified in the processes of differentiated astrocytes, induced by nanotopography [79], is weaker when compared to polygonal astrocytes, which have the shape characteristic of undifferentiated astrocytes grown in the absence of neurons [80–84]. Surprisingly, astrocytes co-cultured with neurons show more frequent dynamics than polygonal cells, and the dynamics were of comparable magnitude. Finally, superresolution imaging by STimulated Emission Depletion (STED) microscopy and analysis with a filament extraction algorithm demonstrates that the specific orientation of actin filament structures occurs only in differentiated cells, indicating that both structure and dynamics carry information about the environment. Overall, our results indicate that actin is dynamic in astrocytes and behaves collectively by clustering into “hotspot” regions that respond in distinct ways to specific chemo-physical stimuli and may drive the homeostatic response of the neural cell network.

## Results

### Primary astrocytes display actin dynamics in response to chemical cues

Our live imaging results demonstrate that astrocytes consistently display actin dynamics, particularly near the boundary of the cell. Representative overlays of fluorescent images are shown in **Fig. 1A** (A1 and inset for representative control timelapse; A2 and inset for representative “triggering” timelapse). In this representation, a perfectly stationary cell would appear all white; the presence of different colors indicates non-overlapping structures at either earlier (more blue) or later (more red) times. To quantify actin dynamics, we employ an optical flow method [85] similar to that used in our previous work [60]. We additionally use shape analysis [57,67,68] to separately consider actin dynamics within the whole cell, boundary region, and bulk region (see **Fig. 1A3-4** for representative analyses and **Fig. 1B** for schematic and *Materials and Methods* for more detail). Still images of actin-GFP and brightfield (separate and merged) are also included (**Fig. 1 – figure supplement 1A**).

**Fig. 1.**
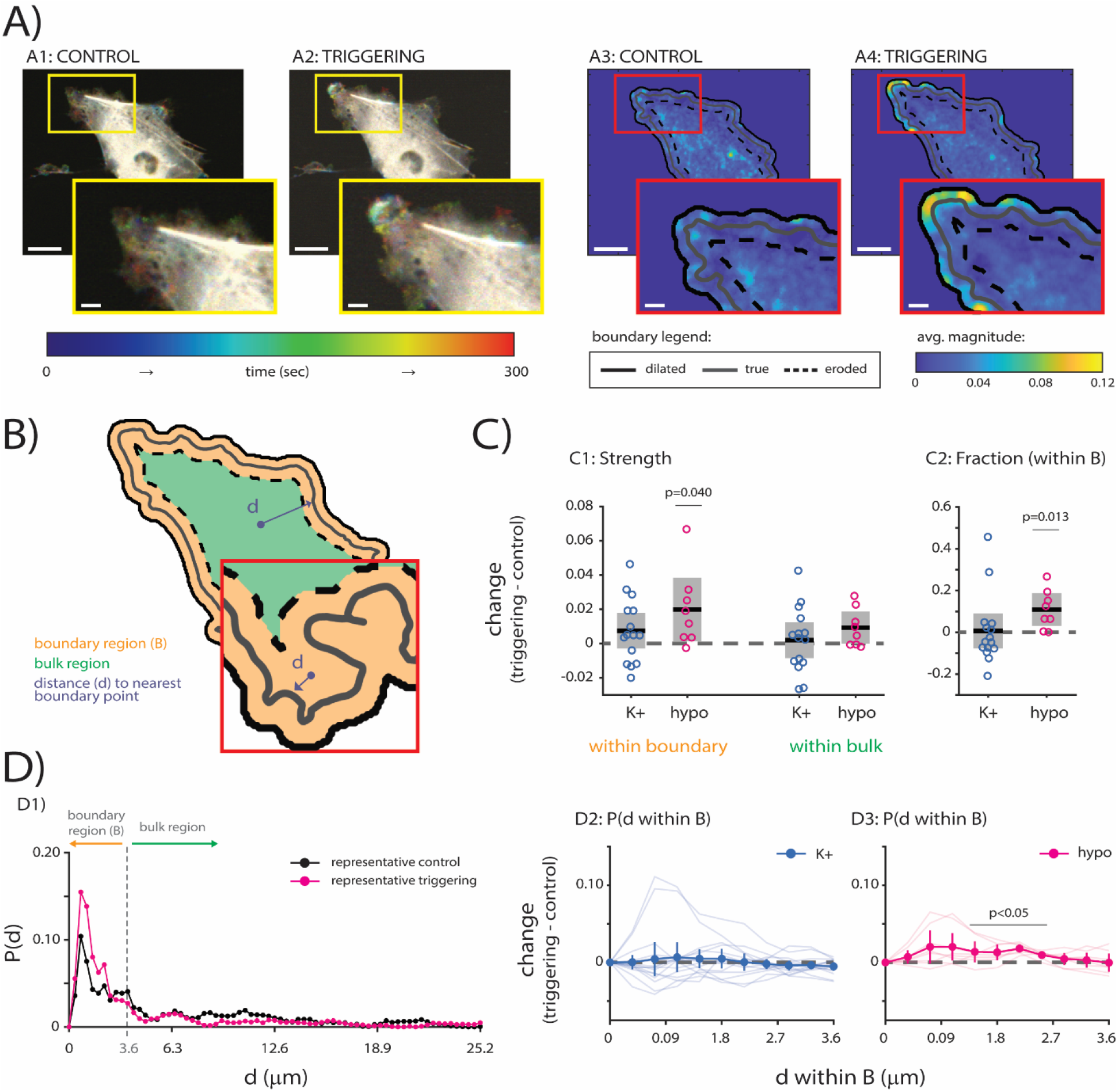
Optical flow analysis reveals actin dynamics in primary astrocytes. **A)** Representative actin dynamics. Actin-GFP timelapse overlay for representative cell, control (A1 and inset) and triggering (A2 and inset) conditions. Colorbar indicates time. Optical flow analysis for representative cell: control (A3 and inset) and triggering (A4 and inset) conditions. Colorbar indicates average optical flow magnitude. **B)** Shape analysis [57,67,68] identifies true boundary (solid grey line). Boundary region (orange) is created by dilating (solid black line) and eroding (dashed black line) true boundary by 10 pixels and is distinct from bulk region (green). **C)** Characterizing actin dynamics. **C1)** Strength of actin dynamics within boundary (p=0.147 for control vs. K+, p=0.040 for control vs. hypo) and within bulk (p=0.696 for control vs. K+, p=0.052 for control vs. hypo). **C2)** Fraction of actin dynamics within boundary (p=0.881 for control vs. K+, p=0.013 for control vs. hypo). **D)** Distance dependence of actin dynamics. **D1)** Probability *P*(*d*) vs. distance from boundary *d* for representative data shown in B. **D2-D3)** Probability within boundary region *P*(*d within B*) vs. distance from true boundary within boundary region *d within B* for all cells for control vs. K+ (D2) and for control vs. hypo (D3; p=0.0499, p=0.019, p<0.001, p<0.001 at 1.44-2.52 μm). P-values determined by paired t-test (C) or by repeated measures ANOVA (D). n=15 paired cells for control vs. K+, and n=8 paired cells for control vs. hypo. Scalebars indicate 20 μm (A1-A4) or 5 μm (insets). Error bars/boxes indicate 95% confidence intervals. Solid black lines and opaque circles indicate mean. Transparent lines in D2-D3 represent individual distance curves. “Control” indicates cells imaged in standard external medium; “triggering” indicates cells imaged in isotonic high potassium solution (“K+”) or hypotonic solution (“hypo”).

To obtain an overall picture of how actin dynamics changes between control and triggering conditions, we first quantified the strength of actin dynamics within specific subcellular regions, specifically the boundary region and bulk region (**Fig. 1C1**). Only triggering with hypotonic causes an increase in the strength of actin dynamics within the boundary region (p=0.040 for control vs. hypotonic via paired t-test). We do not observe significant differences in actin dynamics strength within the boundary after triggering with high K^+^ (p=0.147 for control vs. high K^+^ via paired t-test) or within the bulk for either triggering condition (p=0.696 and p=0.052 for control vs. high K^+^ and for control vs. hypotonic, respectively, via paired t-test). Notably, neither of the triggering conditions changes the probability of actin dynamics within the boundary or bulk regions (**Fig. 1 – figure supplement 1B**).

We next wanted to understand whether actin dynamics becomes more probable within the boundary region after a triggering stimulus. To accomplish this, we calculate the fraction of dynamics within the boundary and find a significant shift in actin dynamics towards the boundary after triggering with hypotonic but not after triggering with high K^+^ (**Fig. 1C2**; p=0.881 and p=0.013 for control vs. high K^+^ and for control vs. hypotonic, respectively, as determined by paired t-test).

To delve into distance dependence of actin dynamics more closely, we quantify the distance from instances of actin dynamics to the cell boundary (true boundary shown in **Fig. 1B**). Curves of the representative cells shown in **Fig. 1A** are shown in **Fig. 1D1** with the boundary and bulk regions labeled. Since we are most interested in the high actin activity near the boundary, we compare how the probability of actin dynamics changes over distance when comparing the two triggering conditions to their matched controls. High K^+^ cells display no difference in actin dynamics at specific distances within the boundary region (**Fig. 1D2**). In contrast, hypotonic cells are significantly more likely to show actin dynamics within the boundary region compared to matched controls, specifically between 1.44 and 2.52 μm from the boundary (p<0.05 for control vs. hypotonic, for as determined by repeated measures ANOVA; refer to **Fig. 1D3** legend for exact p-values). Taken together with the data from **Fig. 1C2**, we show that a higher probability of actin dynamics at specific distances after hypotonic challenge is indeed accompanied by an overall shift to the boundary region.

These results indicate the following: i) actin dynamics has a higher probability of occurring under hypotonic conditions (**Fig. 1C1**); ii) increased probability of actin dynamics at specific distances is observed when cells are exposed to hypotonic challenge (**Fig. 1D3**); and iii) this increase at specific distances after hypotonic challenge is also reflected in the overall probability of actin dynamics (**Fig. 1 – figure supplement 1B**) and in the shift of actin dynamics to the boundary (**Fig. 1C2**).

Finally, we also used inhibitors of the inwardly rectifying K^+^ channel Kir4.1 (barium [Ba^2+^]) and of gap junctions (carbenoxolone [CBX]), involved respectively in potassium spatial buffering [86,87] and cell volume regulation in astrocytes [88,89], to provide a more mechanistic view of how chemical triggering with high K^+^ and hypotonic media changes actin dynamics. These inhibitors decrease actin dynamics such that robust analysis was not possible (representative fluorescent overlays and analysis provided in **Fig. 1 – figure supplement 2**). Future studies will attempt to study the effects of these inhibitors when actin dynamics are in a more excited state.

### Actin dynamics within astrocytes is clustered into active “hotspot” regions

While characterizing actin dynamics within astrocytes, we noticed that many cells seemed to have active regions near the boundary (**Fig. 2A1**) that we suspected were responsible for most of the actin dynamics. To identify these “hotspot” regions, we apply the image processing techniques of erosion and dilation to identify areas of persistent activity (**Fig. 2A2**). After applying thresholds for activity magnitude and probability and hotspot size, we segment regions that meet these criteria (**Fig. 2A3** and inset).

**Fig. 2.**
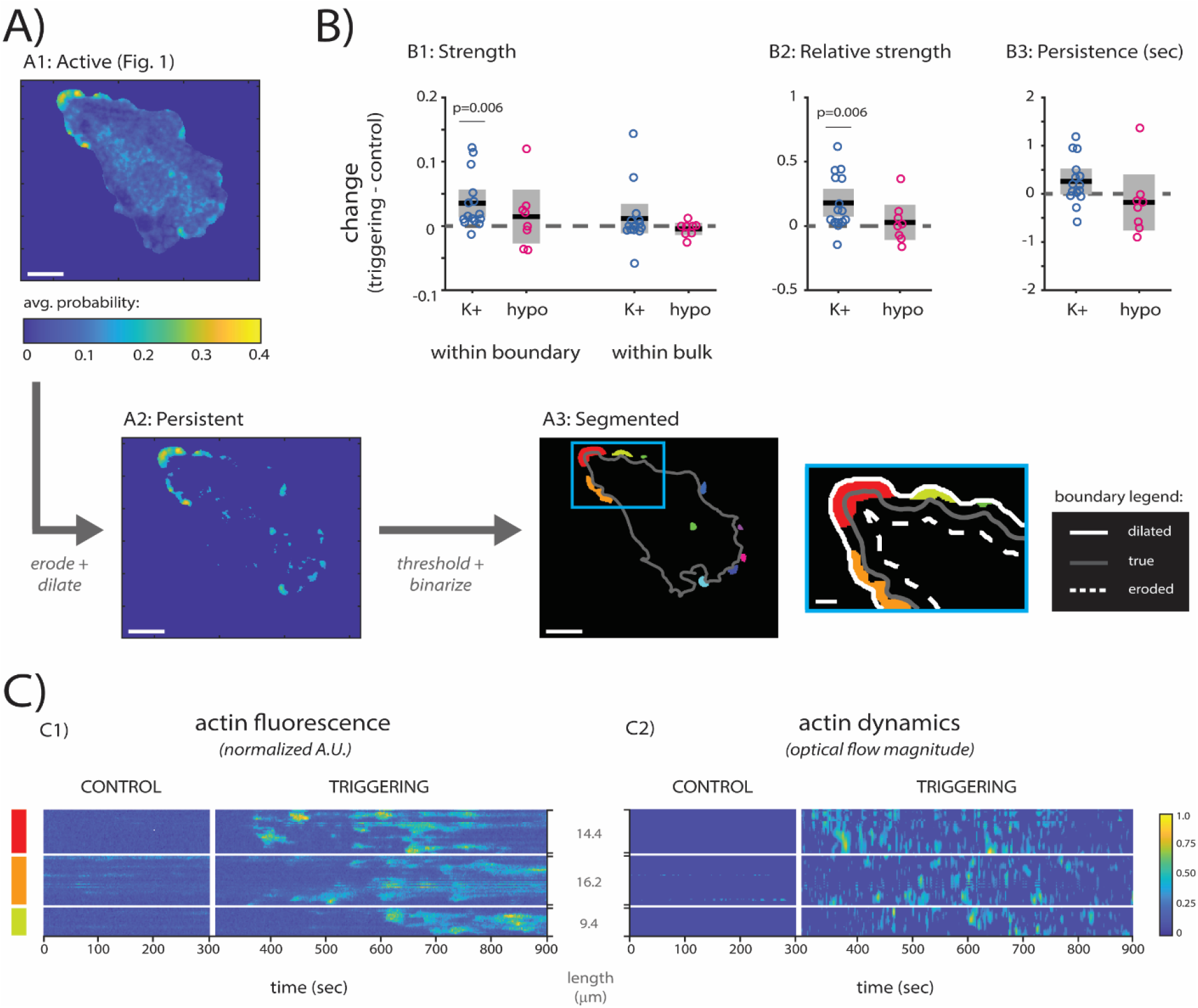
Persistence analysis of actin dynamics reveal “hotspots” of actin activity. **A)** Identification of “hotspot” regions. Optical flow results (A1; from Fig. 1) are eroded and dilated in space and in time (A2) to identify the most persistently active regions. Regions that meet criteria are binarized and segmented (A3, inset A4) for comparison between cells and conditions. **B)** Characterization of hotspot dynamics. **B1)** Average strength of hotspot actin dynamics within boundary (p=0.006 for control vs. K+, p=0.435 for control vs. hypo) and within bulk (p=0.339 for control vs. K+, =0.298 for control vs. hypo). **B2)** Relative strength of actin dynamics (p=0.006 for control vs. K+, p=0.639 for control vs. hypo). **B3)** Average persistence of actin dynamics (p=0.056 for control vs. K+, p=0.491 for control vs. hypo). All p-values determined by paired t-test. Solid black lines indicate the mean. Grey regions indicate 95% confidence interval. **C)** Kymographs of normalized actin fluorescence (C1) and of optical flow magnitude (C2) for three largest hotspot regions from A3-A4. Grey labels between kymographs indicate lengths of regions. Colorbar indicates normalized fluorescence (C1) and optical flow magnitude (C2). n=15 paired cells for control and K+, and n=8 paired cells for control and hypo.

We first compared the strength of actin dynamics within the hotspot regions, considering the boundary and bulk regions separately as before. In contrast to what we found in **Fig. 1C1** (stronger actin dynamics in the boundary region after hypotonic challenge), we find within hotspot regions that actin dynamics is stronger only in cells exposed to high K^+^ and only within the boundary region (**Fig. 2B1**; boundary: p=0.006 for control vs. high K^+^ and p=0.435 for control vs. hypotonic as determined by paired t-test; bulk: p=0.339 for control vs. high K^+^ and p=0.298 for control vs. hypotonic as determined by paired t-test).

We also investigated the relative strength of hotspot dynamics where the average magnitude within hotspots is normalized by the average magnitude of dynamics within the whole cell (**Fig. 2B2**). When comparing this relative strength between cells in triggering conditions and their matched controls, we find that only high K^+^ cells display significantly stronger actin dynamics within hotspot regions compared to the whole cell (p=0.006 for control vs. high K^+^ and p=0.639 for control vs. hypotonic as determined by paired t-test). Finally, we investigated whether actin dynamics in these hotspot regions is more persistent and find that the duration is not significantly different for either high K^+^ cells or hypotonic cells compared to matched controls (**Fig. 2B3**; p=0.056 for control vs. high K^+^ and p=0.491 for control vs. hypotonic as determined by paired t-test).

To complement the results investigating strength and relative strength, we also quantified probability and relative probability for the hotspots after chemical triggers. We find that the probability of actin dynamics in hotspots within the boundary does not change for either stimulus (p=0.969 for control vs. high K^+^ and p=0.557 for control vs. hypotonic as determined by paired t-test), though the probability of dynamics within bulk region hotspots significantly decreases after hypotonic challenge (p=0.847 for control vs. high K^+^ and p=0.027 for control vs. hypotonic as determined by paired t-test; **Fig. 2 – figure supplement 1A**). For both chemical triggers, there is no relative change in probability within hotspots (p=0.215 for control vs. high K^+^ and p=0.462 for control vs. hypotonic as determined by paired t-test; **Fig. 2 – figure supplement 1B**).

To provide a static visualization of actin dynamics within these hotspots, we show traditional actin fluorescence kymographs (**Fig. 2C1**) and novel actin dynamics kymographs (**Fig. 2C2**) for the three largest hotspot regions (red, orange, and yellow) in the representative cell. Both types of kymographs clearly show stronger dynamics in the triggering condition. Moreover, while not evident in the kymograph shown, we also find that cells triggered by high K^+^ show a significant decrease in the number of hotspots (p=0.021 for control vs. high K^+^ and p=0.189 for control vs. hypotonic; **Fig. 2 – figure supplement 1C**) and a concomitant increase in the average area of hotspots (p=0.047 for control vs. high K^+^ and p=0.794 for control vs. hypotonic; **Fig. 2 – figure supplement 1D**), indicating that nearby hotspots may have merged in the high potassium environment. Taken together, these results show that i) hotspot regions respond differently to chemically triggering stimuli than does the subcellular region in which they reside (**Fig. 2B1-B2**) and ii) they are specifically reactive to the high K^+^ chemical stimulus in terms of strength of dynamics and both number and average area of hotspots (**Fig. 2 – figure supplement 1C-D**).

### Astrocytes display distinct actin dynamics in the presence of neurons and nanotopographic cues

To observe actin dynamics in more *in vivo*-like cell culture conditions, we used two different approaches: i) astrocytes plated on a nanostructured surface, with starlike shapes and more realistic molecular and functional features devoted to homeostatic control of potassium concentrations and cell volume [77,79], and ii) co-culture of astrocytes with primary cortical neurons.

For the first approach, we took advantage of the topographic features in hydrotalcyte-like compound (HTlc) films to understand how actin dynamics is altered by mechanical stimuli at the nanometer scale. For the second approach, we transduced primary astrocytes while still in the flask and then added them to cultures of primary neurons to generate a co-culture. The results shown in **Fig. 3** characterize how the actin dynamics of polygonal astrocytes grown on PDL-coated glass (“PDL” or “PDL cells”) differ from differentiated astrocytes grown on the nanotopographic surface HTlc (“HTlc” or “HTlc cells”) and from astrocytes co-cultured with neurons (“Co” or “Co cells”) under control conditions (standard external medium). Representative examples of fluorescence overlays and optical flow analyses for each cell type are shown in **Fig. 3A**. Refer to **Fig. 3 – figure supplement 1A-C** for still images of actin-GFP and brightfield (separate and merged) of astrocytes grown on PDL, on HTlc, and on PDL in the presence of neurons, respectively. Note that the images shown in **Fig. 3 – figure supplement 1A** are the same representative images as those shown in **Fig. 1 – figure supplement 1A**; they are provided again to facilitate comparison between culturing conditions.

**Fig. 3.**
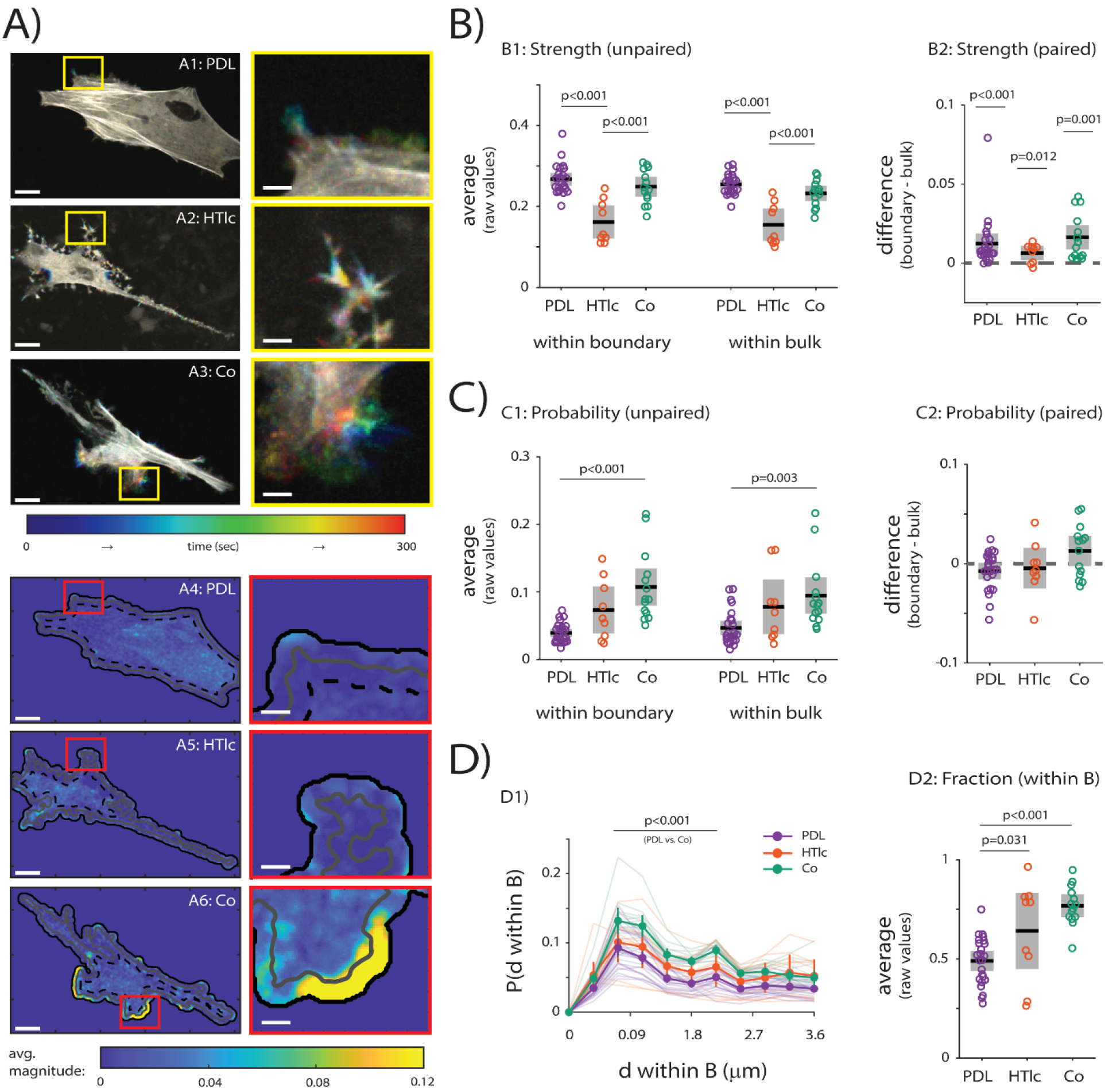
Actin dynamics differs substantially in polygonal astrocytes grown on PDL compared to differentiated astrocytes grown on HTlc and to astrocytes co-cultured with neurons. **A)** Actin-GFP timelapse for representative cells grown on PDL-coated glass (“PDL”; A1 and inset), HTlc films (“HTlc”; A2 and inset), and co-cultured with neurons (“Co”; A3 and inset). Colorbar indicates time. Optical flow analysis for representative cells (PDL in A4 and inset; HTlc in A5 and inset; Co in A6 and inset). Colorbar indicates average optical flow magnitude. **B)** Strength of actin dynamics. **B1)** Strength of actin dynamics within boundary (p<0.001 for PDL vs. HTlc cells, p<0.001 for HTlc vs. Co cells) and within bulk (p<0.001 for PDL vs. HTlc cells, p<0.001 for HTlc vs. Co cells). **B2)** Strength of actin dynamics in boundary vs. bulk for PDL (p<0.001), HTlc (p=0.011), and Co cells (p<0.001). **C)** Probability of actin dynamics. **C1)** Probability of actin dynamics within boundary (p<0.001 PDL vs. Co cells) and within bulk (p=0.003 for PDL vs. Co cells). **C2)** Probability of actin dynamics in boundary vs. bulk for PDL (p=0.072), HTlc (p=0.615), and Co cells (p=0.093). **D)** Distance dependence of actin dynamics. **D1)** Probability within boundary region *P*(*d within B*) vs. distance from true boundary within boundary region *d within B* for all PDL, HTlc, and Co cells (p<0.001 at 0.072-2.16 μm for PDL vs. Co). **D2)** Fraction of actin dynamics within boundary (p=0.031 for PDL vs. HTlc, p<0.001 for PDL vs. Co, and p=0.122 for HTlc vs Co). P-values determined by one-way ANOVA followed by Tukey’s multiple comparisons test (B1, C1, D2), paired t-test (B2, C2), or two-way ANOVA followed by Tukey’s multiple comparisons test (D1). n=24 cells for PDL, n=9 cells for HTlc, and n=14 cells for Co. Scalebars indicate 20 μm (A1-A6) or 5 μm (insets). Error bars/boxes indicate 95% confidence intervals. Solid black lines and opaque circles indicate mean. Transparent lines in D1 represent individual distance curves.

In **Fig. 3B**, we first compared how actin dynamics differs in the boundary and bulk regions for PDL cells, HTlc cells, and Co cells. We find that both PDL cells and Co cells show significantly stronger dynamics within the boundary region and within the bulk region compared to HTlc cells (**Fig. 3B1**; p<0.001 for HTlc vs. PDL and p<0.001 for HTlc vs. Co as determined by one-way ANOVA followed by Tukey’s multiple comparisons test). Surprisingly though, when comparing boundary to bulk within each condition (**Fig. 3B2**), we find that all conditions (PDL, HTlc, and Co) show significantly stronger dynamics at the boundary compared to the bulk (p<0.001 for PDL, and p=0.012 for HTlc, and p=0.001 for Co as determined by paired t-test).

We next compared how likely the different groups of cells are to show actin dynamics in different subcellular regions. Within both the boundary and bulk regions, only Co cells are significantly more likely to show actin dynamics compared to PDL cells (**Fig. 3C1**; p<0.001 for boundary and p=0.003 for bulk as determined by one-way ANOVA followed by Tukey’s multiple comparisons test). The probability of observing dynamics within the boundary and bulk of HTlc cells is not significantly different from either PDL or Co cells (boundary: p=0.051 for PDL vs. HTlc and p=0.082 for HTlc vs Co; bulk: p=0.128 for PDL vs HTlc and p=0.604 for HTlc vs Co; all as determined by one-way ANOVA followed by Tukey’s multiple comparisons test). In this case, there is no significant difference between probability of dynamics within boundary and bulk for any condition (**Fig. 3C2**; p=0.072 for PDL, p=0.615 for HTlc, and p=0.093 for Co as determined by paired t-test).

Finally, we evaluated distance dependence of actin dynamics under each culturing condition in **Fig. 3D**. We show the full distance curves in **Fig. 3 – figure supplement 1D** for the representative cells. When comparing individual distances within boundary region (**Fig. 3D1**), we see that only Co cells are significantly more likely to have actin dynamics than PDL cells at specific distances (p<0.001 for all distances between 0.72 and 2.16 μm as determined by two-way ANOVA followed by Tukey’s multiple comparisons test). Finally, when examining the fraction of events within the boundary (**Fig. 3D2**), we find that PDL cells are less likely to have actin dynamics within the boundary than either HTlc cells and Co cells (p=0.031 and p<0.001 for PDL vs. HTlc and for PDL vs. Co, respectively, as determined by one-way ANOVA followed by Tukey’s multiple comparisons test), with no difference between HTlc and Co cells (p=0.122 as determined by one-way ANOVA followed by Tukey’s multiple comparisons test). Taken together, these results indicate the following: i) HTlc cells show weaker actin dynamics than both PDL and Co cells with no difference in probability (**Fig. 3B1** and **Fig. 3C1**), ii) Co cells show equally strong and more frequent actin dynamics than PDL cells (**Fig. 3B1** and **Fig. 3C1**), and iii) both HTlc and Co cells show a significant shift in dynamics towards the boundary compared to PDL cells (**Fig. 3D2**).

### Astrocytes display hotspot regions in the presence of neurons that are more active than in the absence of neurons

We next sought to compare whether hotspot dynamics differed between astrocytes cultured on PDL-coated glass in the absence of neurons (PDL) and in the presence of neurons (Co). Astrocytes grown and differentiated on HTlc did not display consistent hotspot regions for reliable analysis.

Within hotspots, the strength of actin dynamics within the boundary region is significantly stronger in Co cells compared to PDL cells, whereas we observe no difference between PDL and Co cells within the bulk region (**Fig. 4B1**; p=0.003 for PDL vs. Co within boundary and p=0.063 for PDL vs. Co within bulk as determined by two-sample t-test). Additionally, Co cells also show a significant increase in the strength of hotspot dynamics between the boundary and bulk (**Fig. 4B2**; p=0.075 for PDL and p<0.001 for Co as determined by paired t-tests). The relative strength of hotspots within Co cells is also significantly higher than within PDL cells (**Fig. 4B3**; p=0.009 as determined by two-sample t-test).

**Fig. 4.**
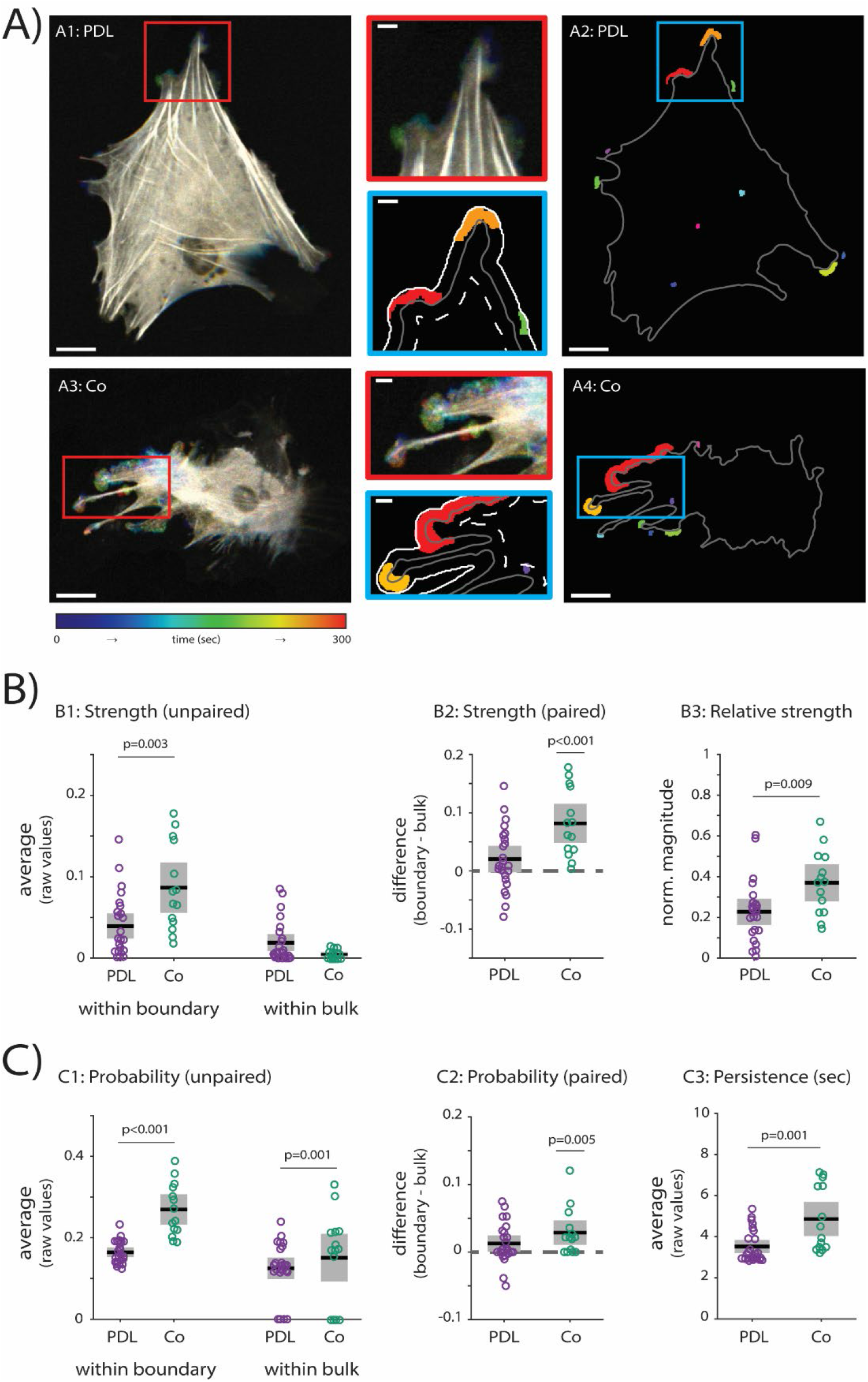
Actin hotspot dynamics are stronger and more frequent when astrocytes are co-cultured with neurons. **A)** Actin-GFP timelapse for representative cells grown on PDL-coated glass (“PDL”; A1 and inset) and co-cultured with neurons (“Co”; A3 and inset). Colorbar indicates time. Persistence analysis for representative cells (PDL in A2 and inset; Co in A4 and inset). Color is used to differentiate hotspot regions. **B)** Strength of actin dynamics within hotspots. **B1)** Strength of hotspot dynamics within boundary (p=0.003 for PDL vs. Co cells) and within bulk (p=0.063 for PDL vs. Co cells). **B2)** Strength of hotspot dynamics in boundary vs. bulk for PDL (p= 0.075) and Co cells (p<0.001). **B3)** Relative strength of hotspot dynamics for PDL vs. Co (p=0.009). **C)** Probability of actin dynamics within hotspots. **C1)** Probability of hotspot dynamics within boundary (p<0.001 for PDL vs. Co cells) and within bulk (p=0.001 for PDL vs. Co cells). **C2)** Strength of actin dynamics in boundary vs. bulk for PDL (p=0.051) and Co cells (p=0.005). **C3)** Persistence of actin dynamics for PDL vs. Co (p=0.001). P-values determined by two-sample t-test (B1, B3, C1, C3) or paired t-test (B2, C2). n=24 cells for PDL, and n=14 cells for Co. Scalebars indicate 20 μm (A1-A4) or 5 μm (insets). Error bars/boxes indicate 95% confidence intervals. Solid black lines and opaque circles indicate mean.

We next considered probability of dynamics within hotspots under different culturing conditions. The probability of hotspot dynamics is significantly higher for Co cells compared to PDL cells, both within the boundary and within the bulk (**Fig. 4C1**; p<0.001 for PDL vs. Co within boundary and p=0.001 for PDL vs. Co within bulk as determined by two-sample t-test). Co cells also show a significant increase in the probability of boundary over bulk hotspot dynamics (**Fig. 4C2**; p=0.051 for PDL and p=0.005 for Co as determined by paired t-tests). Furthermore, the average persistence of dynamics within the hotspots of Co cells is significantly higher than that of PDL cells (**Fig. 4C3**; p=0.001 as determined by two-sample t-test).

Finally, the relative probability of dynamics is lower in Co cells than in PDL cells (**Fig. 4 – figure supplement 1A**; p=0.011 for PDL vs. Co as determined by two-sample t-test), indicating that astrocytes co-cultured with neurons are generally more active than without neurons but that their hotspots are less relatively active compared to astrocytes without neurons. The number of hotspots does not differ between PDL cells and Co cells (**Fig. 4 – figure supplement 1B**; p=0.333 for PDL vs. Co as determined by two-sample t-test), but the average area of hotspots is increased in Co cells (**Fig. 4 – figure supplement 1C**; p=0.021 for PDL vs. Co as determined by two-sample t-test).

Taken together, we have shown that, when compared to primary astrocytes only, astrocytes co-cultured with neurons have hotspot dynamics that i) are stronger and more frequent within the boundary (**Fig. 4B1** and **Fig. 4C1**), ii) are relatively stronger (**Fig. 4B3**) but relatively less frequent (**Fig. 4D1**), and iii) persist for longer (**Fig. 4C3**).

### Astrocytes sense nanotopographic cues through actin orientation near the boundary

Inspired by our analysis of actin dynamics in live astrocytes, we sought to understand i) why certain “hotspots” of actin activity are more prevalent in cells grown on PDL than in cells grown on HTlc and ii) whether the characteristics of actin dynamics are related to the underlying actin structure. To this end, we cultured primary rat astrocytes on PDL-coated glass or on HTlc films and performed STED microscopy on fixed cells stained for F-actin and the intermediate filament protein glial fibrillary acid protein (GFAP). Representative images are shown in **Fig. 5**. HTlc preferentially induces stellate morphology characteristic of differentiated astrocytes, whereas astrocytes cultured on PDL-coated glass display the polygonal shapes characteristic of undifferentiated astrocytes grown in the absence of neurons [12,79–84,90]. We chose not to use astrocytes co-cultured with neurons in this portion of our study because overlapping protrusions would have been visualized via immunostaining due to the density of the co-culture, thus preventing robust single-cell analysis. Cell density did not present a technical or analytical challenge for our dynamics data because transduction efficiency is low enough to obtain single cells in the field of view.

**Fig. 5.**
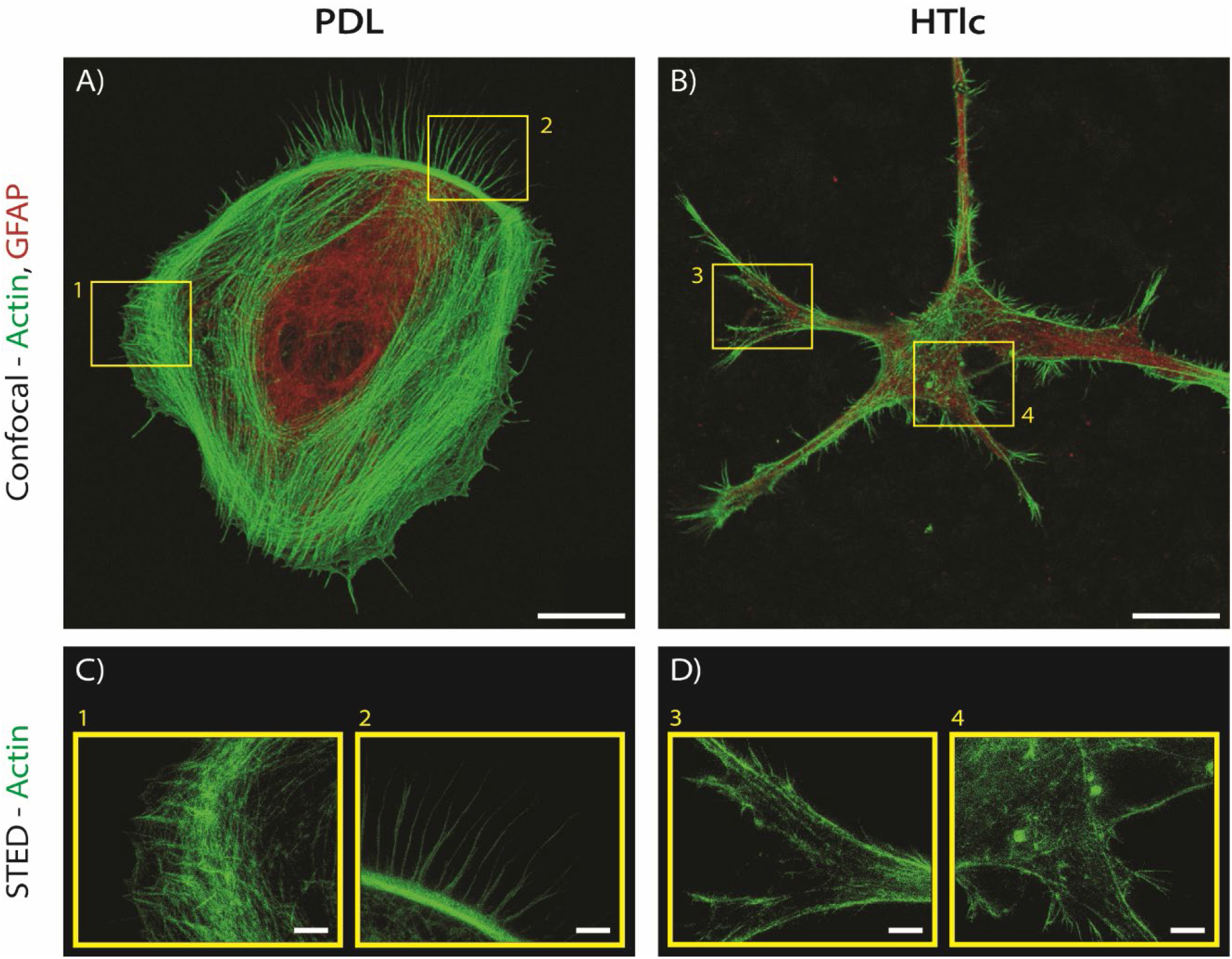
STED microscopy visualizes nanoscale actin structure within primary astrocytes grown on PDL and on HTlc. **A-B)** Confocal microscopy images of representative astrocytes grown on PDL-coated glass (A) or on HTlc films (B) for five days, fixed, and stained for actin (green) and GFAP (red). Scalebars indicate 10 μm. **C-D)** Superresolution images taken via STED microscopy of cells grown on PDL (C) and on HTlc (D) of boxed yellow regions from A and B. Scalebars indicate 2.5 μm.

Confocal images of F-actin labeled astrocytes reveal that there is a marked tendency for cells grown on PDL (**Fig. 5A**) to display a polygonal and dynamic phenotype with distinct leading and trailing edges and several motile structures (i.e. lamellipodia, filopodia, stress fibers; **Fig. 5C**). Conversely, in the “star” shaped cells grown on HTlc (**Fig. 5B**), the same motile elements that would indicate direction of propagation are not clearly visible (**Fig. 5D**). Moreover, the astrocytic actin is seen to “burst” out of the actin cortex in HTlc cells (**Fig. 5D**). This bursting or “flaring” behavior is indicative of a functional microdomain, known to be present in differentiated astrocytes grown on HTlc [79]. Although many regions in the representative HTlc cell display this microdomain morphology, the actin dynamics of astrocytes plated on HTlc is of lower magnitude than that of astrocytes grown on PDL (refer to **Fig. 3B**). Therefore, we chose to investigate how astrocytic actin senses the different mechanical environments presented by PDL-coated glass and HTlc films and analyze the subsequent cytoskeleton remodeling at a nanoscale level via superresolution imaging. We used a method for quantitative, semi-automated analysis of the actin bundles’ preferential orientation [91] to assess the involvement of actin organization in sensing the local mechanical environment around astrocytes.

Image analysis was performed by combining segmentation via an anisotropic, rotating Laplacian of gaussian (LoG) kernel with a hierarchical cluster analysis (see **Fig. 6A** for a schematic overview and *Materials and Methods* for a detailed description). These image processing techniques allow for the analysis of actin angle organization and the determination of fractions of actin roughly parallel versus roughly perpendicular to the leading edge of the cell boundary (**Fig. 6B**). From our clustered data, we determine that PDL astrocytic actin is oriented more parallel relative to the leading edge of the astrocyte, whereas HTlc astrocytic actin is oriented more perpendicular (**Fig. 6C**). The difference seen between the actin of cells grown on PDL versus on HTlc is significant as the cluster algorithm identifies two clusters within the data. Moreover, we see the cluster groups largely correspond with the chosen nanotopography, with an error of seven out of 37 cells: one HTlc cell is misclustered, and six PDL cells are misclustered (i.e., six PDL cells can be found in the cluster of predicted HTlc actin fractions). While the differences observed between astrocytes grown on HTlc and those grown on PDL at a dynamic, micron scale do not necessarily correspond to the differences seen in structure at the nanometer scale, our analytical approach reveals that the underlying nanoscale organization of actin within astrocytes does indeed change when the cells are exposed to different nanotopographic surfaces. Future work will optimize our co-culture approach and determine how the nanoscale structure of astrocytic actin changes when astrocytes are in contact with neurons.

**Fig. 6.**
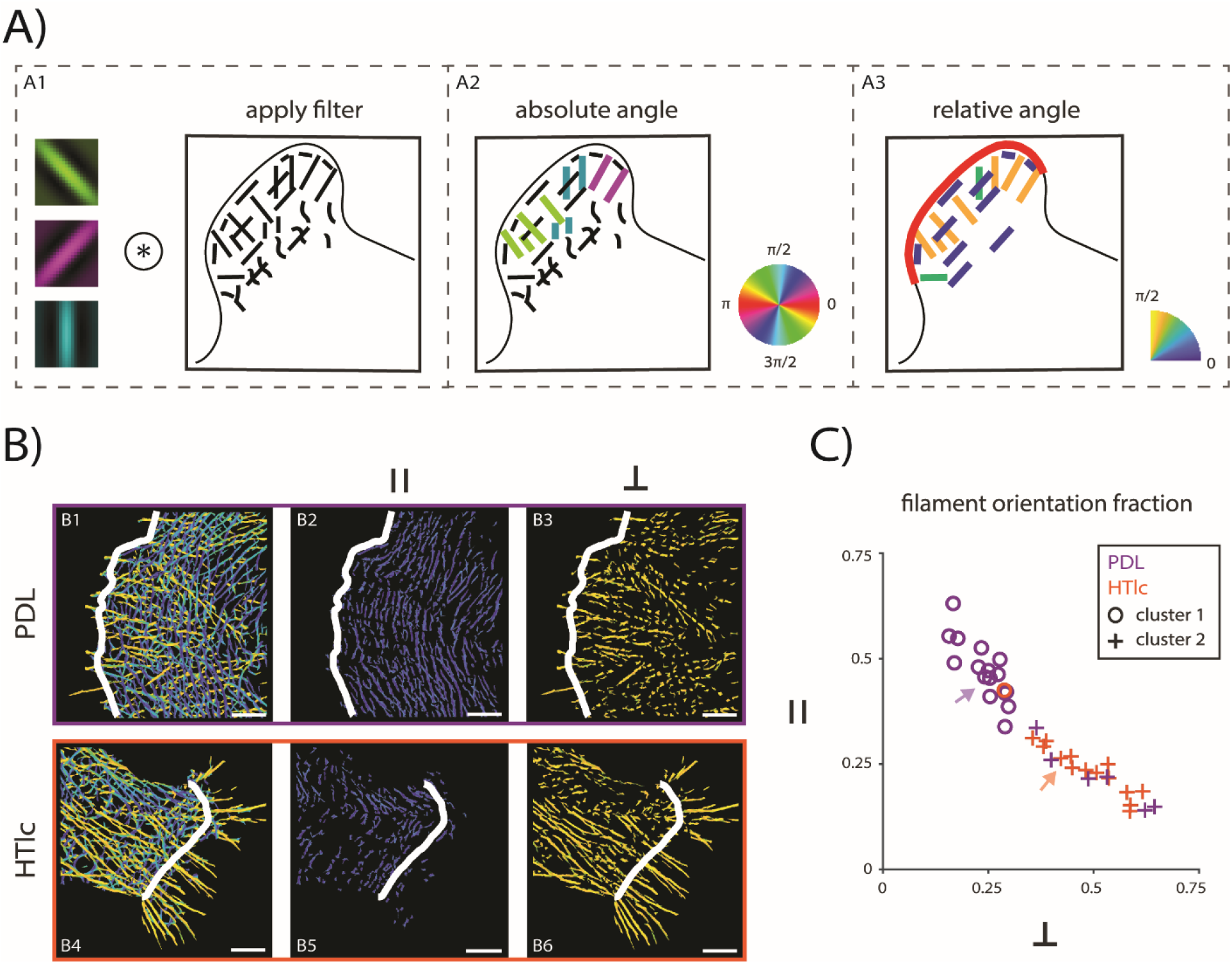
Filtering STED images with a rotating, anisotropic LoG kernel reveals differences in nanoscale actin structure for astrocytes grown on PDL versus on HTlc. **A)** Extraction of relative angle information from actin filaments. **A1)** STED images are convolved with a rotating, anisotropic LoG kernel. **A2)** Filtering extracts actin bundles and a best match angle that is relative to the MATLAB frame (“absolute angle”) with a range of [0, 2π] (shown as full colorwheel). **A3)** A leading edge boundary (red) is drawn manually and is used to transform actin angles to be relative to the closes boundary point (“relative angle”) with a range of [0, π/2] (shown as a quarter colorwheel). **B)** Output from A for representative cells (PDL in B1-B3, HTlc in B4-B6). All relative angles, parallel angles only (range of [0, π/6]), and perpendicular angles only (range of [π/3, π/2]) for representative PDL (B1, B2, B3, respectively) and HTlc (B4, B5, B6, respectively) cells. Solid white lines indicate leading edge boundary. Scalebars indicate 2.5 μm. **C)** Parallel (y-axis) vs. perpendicular (x-axis) fractions of actin detected within PDL (purple) and HTlc (orange) astrocytes. A hierarchical cluster analysis reveals two distinct clusters (cluster 1: open circles, cluster 2: crosses). Arrows indicate representative cells. n=22 cells for PDL, and n=15 cells for HTlc.

## Discussion

We show, for the first time, by means of live timelapse confocal microscopy, that astrocytes respond to alterations in the chemophysical environment with changes in actin dynamics. Furthermore, we demonstrate that actin is spatiotemporally organized into micron scale territories where the dynamics are enhanced depending on the specific chemophysical stress or environment (topography or co-culture). Finally, we show how distinct organization of actin near the boundary may contribute to dynamics.

Our study reveals that actin dynamics occurs persistently and locally, with actin dynamics detectable near the boundary even when the boundary itself does not shift measurably. Hypotonic challenge increases the strength of actin dynamics while high potassium preferentially increases the strength of hotspot dynamics; both changes occur within the boundary region. Furthermore, while astrocytes do not change shape substantially on the short timescales of our experiments, we see small filopodial and lamellopodial structures – the F-actin-rich subcellular compartments of migrating non-neuronal cells [92] – near the cell boundary that appear to “idle” by rapidly extending and retracting, effectively sampling their local environment. It is in these regions, which tend to lack strong stress fibers, that we most often observe actin dynamics. In future experiments, we will quantify how local actin dynamics contribute to changes in astrocyte morphology that occur during differentiation or after injury. Indeed, many studies have extensively demonstrated that, on longer timescales, astrocytes are extremely plastic cells. For example, astrocytes change their morphology by becoming polarized and growing long processes after severe tissue damage caused by traumatic injuries [93–95] and also become hypertrophic and lose their fine processes when they become reactive [19,96].

Surprisingly, dynamics are stronger in astrocytes grown on flat surfaces (PDL-coated glass) and when astrocytes are co-cultured with neurons, clustering into hotspot regions, whereas they are weaker in cells grown on nanostructured surfaces (HTlc films) and do not cluster into large enough regions for hotspot dynamics to be detected with confocal microscopy. Dynamic hotspots in differentiated astrocytes might present as miniature versions of lamellipodia and filopodia, and future work will use higher magnification imaging to determine if this is indeed the case. Importantly, the co-culture results demonstrate that actin dynamics is not highly sensitive to plating density because the co-culture plates were higher in density than the plates with astrocytes only (whether cultured on PDL or HTlc).

Our STED data might also provide the key to understanding why polygonal astrocytes display more robust cytoskeleton dynamics: a more parallel and dispersed organization of actin fibers into a meshwork underneath the cell membrane is likely to confer mechanical plasticity to the plasma membrane compared to differentiated cells grown on HTLc, where actin fibers are packed into “actin rails” and not only maintain the cell shape (especially in the cellular processes) but also might cause membrane stiffness or resistance to motility. In support of our hypothesis, a recent study demonstrated increased stiffness at the leading edge of migrating astrocytes using live STED and atomic force microscopy [97]. We also hypothesize that this distinct actin structure will lead to differences in the interaction with neighboring cells. As stellation enables communication with other cell types, perpendicular actin may ensure processes are directed along the proper axis for communication. Indeed, recent work implicates membrane-bound integrins, specifically astrocytic integrin-engaged Thy1, in interacting with the neuronal cytoskeleton to promote astrocyte-neuron communication [98]. Moreover, our finding that almost all cells cultured on HTlc have actin structures perpendicular to the boundary affirms the ability of HTlc to encourage stellate morphology and differentiation of astrocytes [79]. Our analysis also hints that some cells grown on PDL have actin structures that “look” like the actin structures of stellate, differentiated cells grown on HTlc, consistent with previous observations that growth of astrocytes on PDL delays but does not fully suppress spontaneous or gliotic differentiation that might occur in standard cell culture [48,79,99,100].

Previous studies demonstrated that astrocytes grown on HTlc are more efficient in their homeostatic process. In particular, the expression and function of Kir4.1 and water channel aquaporin 4 (AQP4) as well as the functional response to anisotonic challenge are enhanced in differentiated [77,79] and mature [101–106] cells. A more efficient response to homeostatic challenges due to the higher levels of channel proteins may be related to the weaker actin dynamics observed in differentiated cells compared to polygonal cells (both with and without neurons). While we have not directly investigated this question, a direct relationship between channel function and cytoskeleton dynamics would not be surprising, if we consider that astrocytes may harness the excitable dynamics of actin to codify and convey information coming from the extracellular milieu. Indeed, in fixed cells, it has been shown that the actin cytoskeleton affects AQP4 localization [47,107], but how real time actin dynamics affects the localization and function of AQP4 and other channels, including Kir4.1, remains unknown. To this end, future work will reveal whether the function of potassium and water channels is altered in differentiated compared to polygonal astrocytes after homeostatic challenge and whether any differences in channel localization and function are directly tied to the dynamic response of actin.

Much is known about how co-culturing astrocytes with neurons alters neuronal function [108–111], but how are astrocytes affected by the presence of neurons? Several studies have shown that the presence of neurons and neuronal activity controls gene expression in astrocytes [80,112,113] and affects astrocytic calcium dynamics [114,115]. For example, rat cortical astrocytes express glutamate-aspartate transporter (GLAST), but in the presence of cortical neurons, as we have done here, astrocytes also express glutamate transporter-1 (GLT-1) [80,113]. Neurons have also been shown to enhance astrocytic expression of Cx43, gap junctional communication between astrocytes, and the propagation of calcium waves within the astrocytic network [116]. We hypothesize that actin dynamics in astrocytes co-cultured with neurons is enhanced specifically because actin dynamics contributes to astrocytes’ critical function of facilitating neuronal activity by interacting with neurons and sensing the extracellular environment. Recent work has indicated that actin-dependent morphology changes in PAPs are not only critical for interacting with neurons but may also enhance spine stability [117]. Future work will probe more deeply into how astrocytic actin dynamics affect and may sense the functional dynamics of astrocyte-neuron networks. Indeed, our recent work has shown that actin waves in *D. discoideum* can sense electric fields [64]. We will, in the future, investigate whether actin dynamics in astrocytes can also sense and respond to the electrical activity of neurons.

These questions remain to be elucidated, and studies in microglia [53] – which demonstrated that actin dynamics is critical for their principal function, surveillance – indicate that actin dynamics also plays an important role *in vivo* in neural cells. We propose that actin dynamics in astrocytes is critical for astrocytes’ principal functions: maintaining homeostasis and modulating neuronal communication. Finally, given that cytoskeletal alterations are reported in a variety of neurological conditions characterized by a gliotic state [118], we expect investigations of cytoskeletal dynamics in defined animal models to be an important area of exploration to validate the pathophysiological relevance [119] of our findings.

Moreover, while we know that the astrocyte-neuron interactions will happen on a spatial scale much smaller than the dynamics observed here, our hypothesis is supported by a recent study highlighting the importance of mechanical signaling for synaptic activity [54]. We therefore intuit that astrocytic actin dynamics may serve as a mechanical regulator of neuronal activity.

## Materials and Methods

### Rat cortical astrocyte culture preparation, maintenance, plating, and differentiation

#### Cell culture

Primary astrocytes were obtained from Sprague Dawley rats housed at the University of Maryland (in concordance with the recommendations of and approval by the University of Maryland Institutional Animal Care and Use Committee; protocols R-JAN-18-05 and R-FEB-21-04) or from Wistar rats housed at the University of Bologna (in concordance with the Italian and European law of protection of laboratory animals and the approval of the local bioethical committee, under the supervision of the veterinary commission for animal care and comfort of the University of Bologna and approved protocol from Italian Ministry of Health; ethical protocol number ID 360/2017 PR, released on May 2017, valid for 5 years).

Primary cultures of astrocytes were prepared as described by McCarthy and de Vellis [120] from newborn rat pups between postnatal days 1 and 2 [14,121,90]. Briefly, neonatal cerebral cortices devoid of meninges were gently triturated, filtered with a 70 μm cell strainer (**cat. no. 22-363-548**; Fisher Scientific, MA, USA), and plated in T25 cell culture flasks (**cat. no. 229331**; CELLTREAT Scientific Products, MA, USA) containing Dulbecco’s Modified Eagle Medium with GlutaMAX and high glucose (DMEM; **cat. no. 10-566-024**; Fisher Scientific, MA, USA) supplemented with 15% fetal bovine serum (FBS; **cat. no. 100-106**; Gemini Bio-Products, CA, USA) and penicillin-streptomycin (P/S) at 100 U/mL and 100 μg/mL, respectively (P/S; **cat. no. 15070063**; Thermo Fisher Scientific, MA, USA). Flasks were maintained in an incubator at 37 °C, 5% CO_2_, and proper humidity levels for two weeks. During this period, we replaced cell medium every two days, and flasks were gently shaken when necessary to remove undesired microglial cells. When confluence was reached, astrocytes were dispersed using trypsin-EDTA 0.25% (**cat. no. 25200056**, Thermo Fisher Scientific, MA, USA), and the cell suspension was re-plated on coverslips or in imaging dishes functionalized with Poly-D-lysine hydrobromide (PDL; **cat. no. P7886**; Sigma-Aldrich, MO, USA) as previously described [79,121–124] or with HTlc as previously described [79,125]. Cells were plated at a density of 5-10×10^3^ per imaging dish and maintained in DMEM containing 10% FBS and 1% P/S.

For co-culture experiments, astrocytes were prepared as described above. At one to two weeks after the astrocyte dissection, neurons were prepared as previously described from rats at embryonic day of gestation 18 (E18) [126–131]. Briefly, embryos were removed via Caesarian section and decapitated. Cerebral hemispheres were separated, and meninges were removed. The hippocampi and cortices were dissected, and cortices were combined and manually triturated. Cortical neurons were plated in imaging dishes coated with PDL at a density of 5×10^5^ cells/dish and were maintained in NbActiv4 medium (**cat. no. Nb4-500**, BrainBits, TN, USA) with 1% P/S. When astrocyte flasks reached confluence, astrocytes were added to the neuronal culture at a density of 1-2×10^5^ cells/dish, and the media was changed to DMEM with 10% FBS and 1% P/S.

As can be seen in **Fig. 1 – figure supplement 1A** and **Fig. 3 – figure supplement 1A-C**, plating density was chosen to ensure that cultures were not confluent (which would prevent reliable transduction) and also that astrocytes were not isolated (which could affect dynamics).

#### PDL and HTlc film preparation

Plastic dishes with glass inserts (**cat. no. P35G-1.0-20-C**, MatTek, MA, USA) or glass coverslips were functionalized with PDL to serve as a control surface or with Zn-Al hydrotalcite nanoparticles (HTlc) to induce morphological differentiation of astrocytes. For live imaging, PDL was used at a working concentration of 0.1 mg/mL. For STED imaging, PDL was used at a working concentration of 0.01 mg/mL. In both cases, coverslips were coated with PDL for 20 minutes at room temperature and washed three times with approximately 500 μL of sterile H2O. Coverslips or dishes were functionalized with HTlc by adding the HTlc suspension drop by drop on the coverslips and allowed to dry overnight. Once dried, the HTlc coverslips were sterilized by UV for approximately 30 min. For live imaging, astrocytes grown on PDL and HTlc were cultured for three to seven days after replating prior to imaging, and co-cultured astrocytes were cultured for two to three days prior to imaging. For co-culture imaging, neurons were between 4 and 18 days *in vitro*. For STED imaging, astrocytes were cultured for five days after replating prior to fixation.

#### Transduction of cells with actin-GFP

To visualize actin dynamics, we chose to transduce astrocytes with CellLight Actin-GFP, BacMam 2.0 (**cat. no. C10506**; ThermoFisher Scientific, MA, USA) because it has minimal toxicity and sparsely labels cells, which is ideal for single cell analysis. For PDL and HTlc cells, transduction occurred in the imaging dish at a concentration of 100 particles per cell (PPC). At 48 hr prior to live imaging, 5-10 μL (depending on the plating density) of the reagent was added to the cell culture media. A full media change was performed 16-20 hr after transduction, and imaging was performed 24-48 hr after the full media change. For co-cultured astrocytes, transduction occurred in the flask at a concentration of 10 PPC. At 24 hr prior to plating, 100 μL of the reagent was added to the cell culture medium within the flask. Plating was performed 16-20 hr after transduction. Imaging was performed 48-72 hr after plating.

#### Live confocal imaging of transduced astrocytes

Live imaging timelapses were acquired at the University of Maryland CMNS Imaging Core using a PerkinElmer spinning disk confocal microscope with an oil immersion 40x objective (1.30 NA; 0.36 μm/pixel) and under temperature, CO_2_, and humidity control. The microscope was equipped with a Hamamatsu ImagEM X2 EM-CCD camera (C9100-23B), which recorded 16 bit images. Data acquisition was performed with PerkinElmer’s Volocity software (version 6.4.0). A subset of timelapses were taken at the CNR on an inverted confocal microscope (Crisel Instruments) using a 40x water immersion objective (0.16 μm/pixel).

Transduced astrocytes were identified by a positive signal in the 488 nm laser channel. After imaging parameters were set, the media was changed from growth media to standard external solution (control) followed by one of the triggering conditions (high potassium solution or hypotonic solution). Single-plane images were acquired every 2 sec in the 488 channel. A consistent z-plane was assured with Perfect Focus (Nikon PFS).

#### Immunostaining and superresolution imaging of fixed astrocytes

##### Antibodies

Mouse monoclonal Anti-Glial Fibrillary Acidic Protein (GFAP) antibody (diluted 1:200; **cat. no. G3893**; Sigma-Aldrich, MO, USA) and Goat anti-Mouse IgG (H+L) Highly Cross-Adsorbed Secondary Antibody, Alexa Fluor 594 (diluted 1:300; **cat. no. A-11032**; Thermo Fisher Scientific, MA, USA) were used as primary and secondary antibody, respectively, for GFAP immunostaining. Alexa Fluor 488 Phalloidin (diluted 1:500; **cat. no. A12379**; Thermo Fisher Scientific, MA, USA) was used for direct labelling of actin.

##### STED immunostaining

Immunofluorescence for gated STED (gSTED) was performed as indicated in the Quick Guide to the STED Sample Preparation (www.leica-microsystems.com), with some slight adjustments as previously reported [132,133]. Astrocytes were plated on the PDL and HTlc-treated coverslips and cultured for five days. On the fifth day of culture, cells were fixed with 2% paraformaldehyde (**cat. no. P6148**; Sigma-Aldrich, MO, USA) in phosphate buffered saline (PBS) for 10 min, washed with PBS, and permeabilized with 0.3% Triton X-100 (**cat. no. T8787**; Sigma-Aldrich, MO, USA) for 10 min. After blocking with 0.1% gelatin in PBS, fixed astrocytes were incubated with GFAP-primary antibody for 1 hr at room temperature. Cells were then rinsed with 0.1% gelatin-PBS and co-incubated with Alexa Fluor 488 Phalloidin and Alexa Fluor 594-conjugated secondary antibody for 1 hr at room temperature. After washing with PBS, coverslips were mounted on microscope slides by using the ProLong Glass Antifade Mountant (**cat. no. P36980**; Thermo Fisher Scientific, MA, USA) without DAPI, as indicated in Leica official guide, and imaged with both confocal and STED microscopy.

##### STED imaging

Confocal and STED images of fixed astrocytes grown on PDL and HTlc were acquired using a Leica TCS SP8 3X microscope, provided with AOTF and AOBS, white light laser (WLL), Hybrid Detectors (HyD), and two STED lasers (592 nm, 660 nm) [133]. A Leica HC PL APO 100x/1.40 NA Oil STED White objective and Type F Immersion liquid with a refractive index of 1.5 were used. Before starting imaging, the excitation and the doughnut-shaped STED beams were switched on (WLL set laser power= 70%; STED-592 nm set laser power = 98%), aligned, and allowed to reach operating temperature. The beam alignment was repeated whenever necessary. Excitation of the Alexa Fluor 488 dye was achieved using a continuous-wave 488 nm laser line (NKT Photonics supercontinuum laser). For superresolution imaging of actin, g-STED was performed using the continuous wave 592 nm-emitting STED fiber laser. More detailed acquisition settings are reported in **Table 1**. All confocal and STED images were acquired at a set room temperature of 20 °C under constant Acquisition Mode settings (Format scanning resolution: 1024×1024 pixels; Scan Speed: 100 Hz).

**Table 1:** Imaging parameters. Parameters used for STED and confocal imaging of fixed astrocytes on PDL and HTlc.

|  | Parameter | Actin | GFAP |
| --- | --- | --- | --- |
| Confocal | WLL (488 nm) intensity, % | 10 | 10 |
|  | Gain | 25 | 25 |
| STED | WLL (488 nm) intensity, % | 40 |  |
|  | STED (592 nm) intensity, % | 97 |  |
|  | Gain | 200 |  |

##### Criteria for acquisitions of ROIs

Single or few and well-spaced cells were preferentially chosen for STED imaging to ensure no overlap of actin structure belonging to different cells. Actin and GFAP images of the whole cell were first acquired by confocal microscopy at an original magnification of 100x. Then, multiple random regions of interest (ROIs) (~20×20 μm sized) were selected and imaged using an extra optical zoom (from 5 to 7) by confocal and STED microscopy. Three independent experiments were conducted.

#### Synthesis of ZnAl-HTlc nanoparticles and film preparation

Colloidal aqueous dispersion of ZnAl-HTlc nanoparticles having the formula [Zn_0.72_Al_0.28_(OH)_2_] Br_0.28_ 0.69 H_2_O were prepared by the double-microemulsion technique previously described [79]. For the film preparation, a 125 or 160 μL aliquot of ZnAl-HTlc colloidal dispersion at a concentration of 1.2 mg/mL was dropped onto 15 or 19 mm diameter glass coverslips and successively dried for 4 hr in a sterile hood. The obtained ZnAl-HTlc films were used to culture astrocytes and as substrates for live dynamics imaging and fixed STED imaging as described above. In this manuscript, the ZnAl-HTlc films are referred to simply as HTlc.

#### Solutions and chemicals

All salts and chemicals employed for the investigations were of the highest purity grade. For live imaging experiments, the standard external solution (“control”) was the following (mM): 140 NaCl, 4 KCl, 2 MgCl_2_, 2 CaCl_2_, 10 HEPES, 5 glucose, pH 7.4 with NaOH and osmolarity adjusted to ~315 mOsm with mannitol. When using external solutions with different ionic compositions, such as the high potassium solution (“high K^+^”, [40 Mm]), salts were replaced equimolarly. The hypotonic extracellular solution (“hypotonic”) of 260 mOsm/L was obtained by omitting mannitol in the control solution. When imaging, media was changed carefully to prevent accidental, mechanical stimulation of cells with the pipette. Moreover, we controlled for any effects that changing media might cause by changing media prior to imaging under control conditions (from growth media to the standard external solution) and under triggering conditions (from the standard external solution to high K^+^ or hypotonic medium).

#### Analysis of live imaging data

##### Pre-processing of timelapses

All timelapses were smoothed in time using a Simoncelli smoothing filter [134]. Prior to smoothing, any timelapses that showed noticeable jitter or drift due to thermal gradients were processed through a jitter correction algorithm using discrete Fourier transform (DFT) registration and an upsampling factor of 100 (using the “dftregistration” code package from [135]). For timelapses where drift was inconsistent and unable to be fully corrected with the DFT method, an additional correction using intensity-based image registration (via the MATLAB functions “imregconfig” and “imregister”) was applied using the average image of the DFT corrected timelapse as the fixed image.

##### Shape analysis and identification of bulk and boundary regions

We used shape analysis to identify cell boundaries and subsequently to be able to compare actin dynamics within different regions of astrocytes. Using a snake algorithm from our previous work [57,67,68], we analyzed the average fluorescent image of each cell to generate the “true boundary” (**see Fig. 1A-B**). Prior to shape analysis, images were smoothed (“imgaussfilt” in MATLAB with a standard deviation of 1.0) and subsequently sharpened (“imsharpen” in MATLAB with a radius of 2.5 and amount of 1.0). Alpha and beta parameters for the snake algorithm (tension and rigidity, respectively) were 0.20 and 0.25, respectively. We found that this true boundary tended to capture the most prominent morphological features of the cell but often excluded regions that were only transiently dynamic. To mitigate this challenge, we eroded or dilated the true boundary using a spherical structural element with a radius of 10 pixels. By subtracting the eroded cell region from the dilated cell region, we identified a boundary region. We considered the eroded cell mask to be the bulk region.

##### Optical flow analysis of actin dynamics

Optical flow analysis similar to that used in [60] was performed on smoothed and jitter-corrected timelapses in Python 3.8. The optical flow weight matrix was a 19×19 pixel Gaussian with a standard deviation of 3 pixels (1.1 μm) for all timelapses taken at UMD. For a small subset of HTlc timelapses taken at the CNR, the optical flow weight matrix was a 41×41 pixel Gaussian with a standard deviation of 6.75 pixels (1.1 μm). The truncation value for the Gaussian kernel was consistently set to 3. Because actin dynamics in astrocytes is less prominent than in migrating cells usually studied by our lab [56,60], we only considered optical flow vectors with the highest 1% of magnitude within the dilated cell boundary that also passed our reliability criteria (at least 33% of the mean reliability value for that timelapse). These strict criteria ensured that we were only considering dynamic actin events that are large and reliable compared to background noise.

##### Segmentation of active regions (“hotspots”) of actin dynamics

To identify regions with the most actin activity, we further analyzed our thresholded data by eroding and dilating in the x, y, and time dimensions using an elongated cube with dimensions of [3,3,2]. To ensure that we were not including regions of low activity, we set an event probability threshold of 0.10, meaning that a region had to be active at least 10% of the time to be considered a “hotspot”. Furthermore, we imposed a size constraint of 20 pixels (including the time dimension) to ensure that any active regions did not just barely meet the volume require of 3×3×2. Since the dynamics within HTlc cells are on a smaller scale, the hotspot analysis did not reveal any regions that were clearly active.

#### Analysis of STED images of fixed astrocytes

##### Laplacian of Gaussian (LoG) filtering to segment actin

To characterize the organization of the actin meshwork in STED images of fixed cells, as shown in **Fig. 5**, we performed several image processing techniques using custom-written algorithms in MATLAB (Mathworks). Prior to processing, all STED images are adjusted using a contrast-limited adaptive histogram equalization (“adjusthisteq” function in MATLAB’s proprietary API (https://www.mathworks.com/help/images/adaptive-histogram-equalization.html). The resolution of the STED images is 16.18pix/nm. Next, each image is convolved with an anisotropic, rotating Laplacian of Gaussian (LoG) kernel. The exact kernel parameters were determined through trial and error to approximately match the cylindrical shape of actin as visualized by Phalloidin-488 staining. The number of angles through which the LoG kernel is rotated was chosen to balance computational time and segmentation accuracy. For each pixel, the best match angle is chosen via the maximum value resulting from convolving that pixel with all rotations of the filter, and a threshold is applied to ensure a high-quality match. This processing results in a filtered image that highlights both the actin organization and the angle associated with each filament extracted from kernel (see **Fig. 5A1**). The angles associated with the extracted actin are defined in terms of the Cartesian coordinate system, which is not biologically meaningful. To establish the angle at which the actin organizes relative to the cellular boundary, we manually draw a boundary for each cell. The number of pixels per boundary varies on a cell-to-cell basis. For each pixel, we derive the angle of that pixel’s actin relative to the closest boundary point. This transformation generates angles in the range of 0 to π/2. We then apply an additional threshold at an arbitrary value to only segment the longest actin (**Fig. 5A2**); doing so greatly reduces the noise in our processed data. We analyzed at least 15 individual cells per nanotopographic surface, and the morphologies of the imaged areas varied from cell to cell. Thus, the boundaries are generated to normalize the distributions of the relative angles across the data sets. We noticed that for the STED images taken of astrocytes grown on HTlc, there is more total boundary per unit-area. Thus, the boundaries from which the relative angles are calculated are drawn to coincide with only the leading edge of the cell (see **Fig. 5B** for representative examples).

##### Cluster analysis of relative angles

To understand how astrocytic actin organization differs when the cells are grown on different nanotopographies, we use a hierarchical cluster analysis from MATLAB’s proprietary API (https://www.mathworks.com/help/stats/cluster-analysis.html). From the relative angle distributions across both PDL and HTlc (22 individual cells and 15 individual cells, respectively), we group the distributions into “parallel” (angles between 0 and π/6) and “perpendicular” (angles between π/3 and π/2), as shown in **Fig. 5B**. As inputs to the clustering algorithm, we use the fractions of “parallel” and “perpendicular” actin of the 37 individual cells but do not include information about the corresponding nanotopographic surface. We then use the “pdist,” “linkage,” “cophenet,” and “linkage” functions (while optimizing the cophenetic coefficient of our ‘distance’ and ‘linkage’ parameters) to generate predictive clusters of the input data.

#### Data representation and statistics

All data were plotted and all statistical tests performed in MATLAB 2021a. For data showing differences between control and triggering or between boundary and bulk regions, paired t-tests were used (“ttest” function in MATLAB). For data in **Fig. 3** comparing PDL to HTlc to Co, one-way analysis of variance (ANOVA) tests were used (“anova1” function). For data in **Fig. 4** comparing PDL to CO, unpaired two-sample t-tests were used (“ttest2” function). When analyzing probability of actin dynamics vs. distance, ANOVA tests were used as follows: repeated measures ANOVA for paired data in **Fig. 1** (“fitrm” and “ranova” functions), and two-way ANOVA to compare PDL to HTlc to Co in **Fig. 3** (“anovan” function). For all ANOVA tests, Tukey’s multiple comparisons test (“multcompare” function) was used as the post test. All errorbars indicate 95% confidence interval, and the significance level was set at α=0.05. Adobe Illustrator (version 25.0.1) was used for final assembly of all figures.

## Acknowledgements

The authors would like to thank the University of Maryland Imaging Incubator Core Facility for providing and maintaining the systems used in collecting images for this work. They also thank Q. Yang and L. J. Campanello for helpful discussions related to analysis.

## Funding

This work was supported by the following grants:

US Air Force Office of Scientific Research MURI grant FA9550-16-1-0052 to WL
US Air Force Office of Scientific Research grant FA9550-19-1-0370 (AstroDyn) to WL VB GPN KMO
US Air Force Office of Scientific Research FA9550-20-1-0386 to (ASTROLIGHT) to VB
US Air Force Office of Scientific Research FA9550-20-1-0324 to GPN
HORIZON EUROPE SEEDS INTERGLIO (S08) funded by the University of Bari Aldo Moro to GPN
NIH R21NS116892-01 to GPN
US Air Force Office of Scientific Research FA9550-18-1-0255 (3D Neuroglia) to VB
COMBINE fellowship to NJM

## Author contributions

Conceptualization: VB WL

Surface preparation: ES TP

Experiments: KMO ES BB MGM

Analysis: KMO NJM SP

Interpretation: KMO ES BB NJM MGM GPN VB WL

Supervision: GPN VB WL

Writing—original draft: KMO ES BB NJM GPN VB WL

Writing—review & editing: KMO ES BB NJM MGM SP TP RZ GPN VB WL

## Conflict of interest

The authors have no competing interests to declare.

## Figure Supplements

**Fig. 1 – figure supplement 1:**
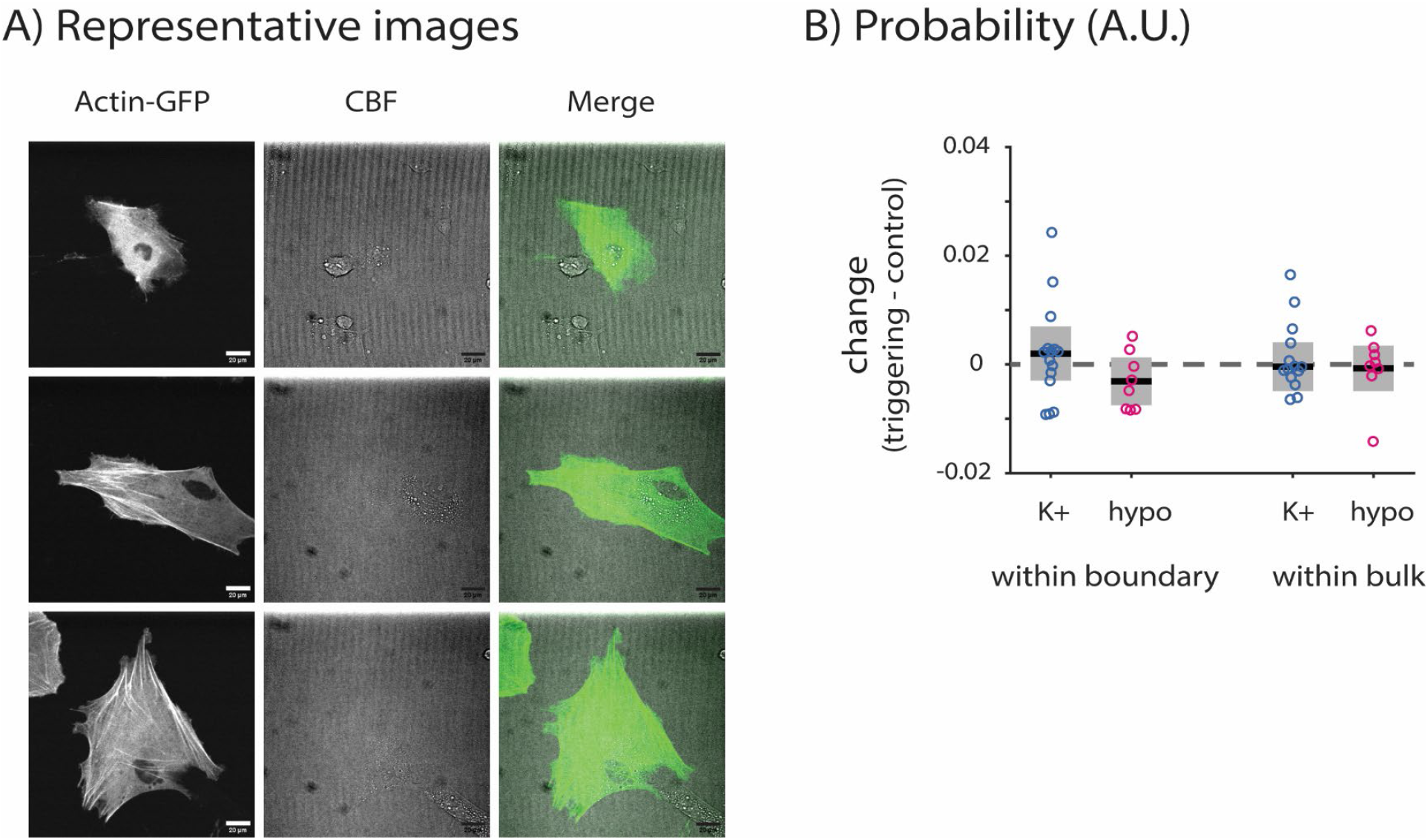
Representative images of primary astrocytes and supplementary data. **A)** Three representative images of primary astrocytes grown on PDL-coated glass (PDL). **B)** Probability of actin dynamics within boundary (p=0.402 for control vs. K+, p=0.138 for control vs. hypo) and within bulk (p=0.854 for control vs. K+, p=0.742 for control vs. hypo). P-values determined by paired t-test. n=15 paired cells for control and K+, and n=8 paired cells for control and hypo. Grey boxes indicate 95% confidence intervals. Solid black lines indicate mean.

**Fig. 1 – figure supplement 2:**
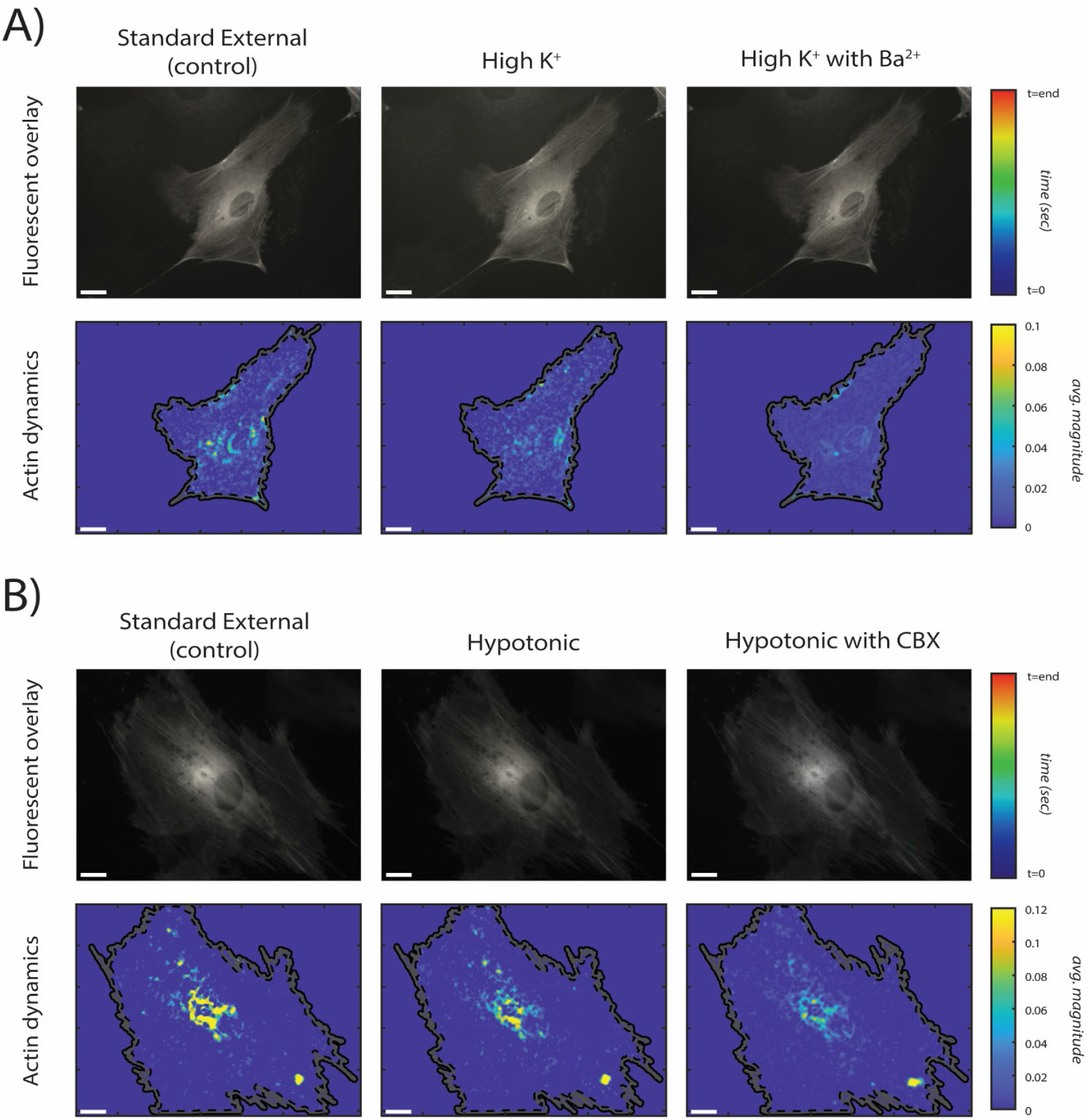
Representative images and analysis of actin dynamics after pharmacologic perturbations. **A)** Representative actin-GFP timelapse overlays (top row) and quantified actin dynamics (bottom row) after triggering with high K^+^ and subsequently blocking potassium channels with barium (Ba^2+^). Left to right: representative astrocyte in standard external (control), high K^+^, and high K^+^ with Ba^2+^. **B)** Representative actin-GFP timelapse overlays (top row) and quantified actin dynamics (bottom row) after triggering with hypotonic and subsequently blocking gap junctions with carbenoxolone (CBX). Left to right: representative astrocyte in standard external (control), hypotonic, and hypotonic with CBX. Top rows: colorbar indicates time. Bottom rows: colorbar indicates average optical flow magnitude. Scalebars indicate 20 μm.

**Fig. 2 – figure supplement 1:**
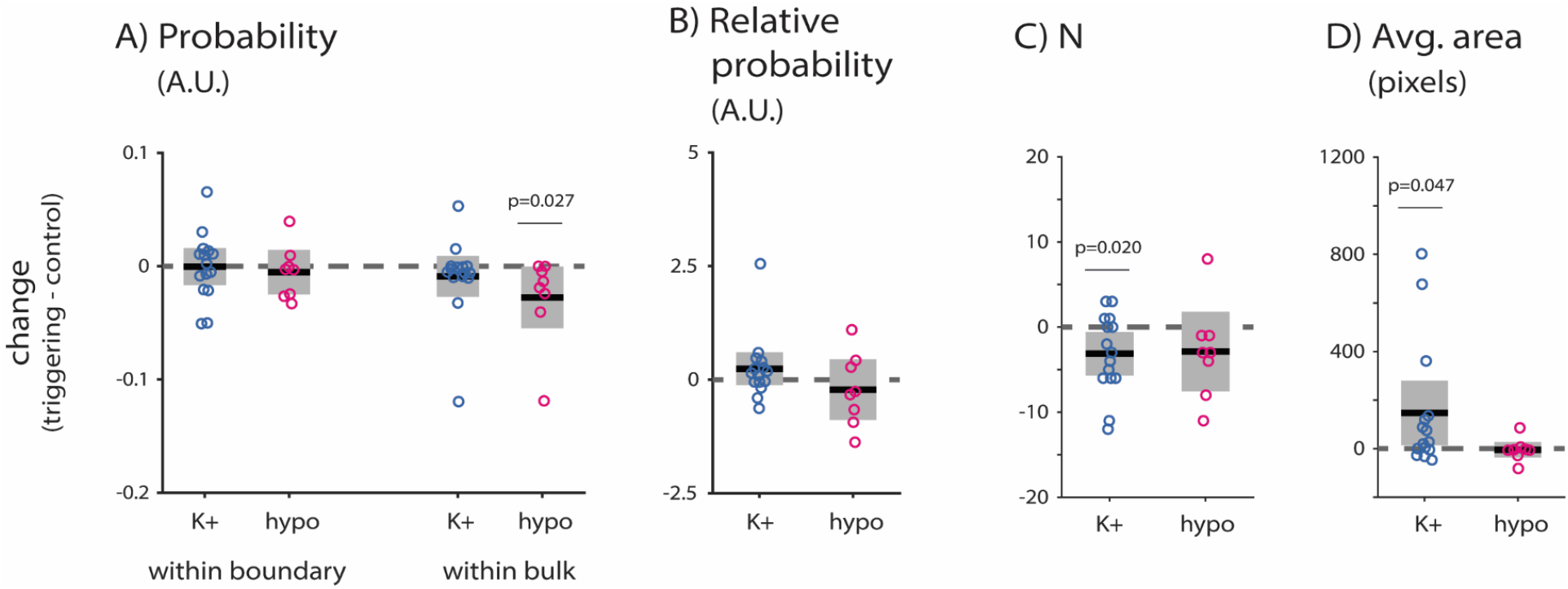
Additional parameters characterizing differences in actin dynamics within hotspot regions after chemical triggering. **A)** Probability of hotspot dynamics within boundary (p=0.969 for control vs. K+, p=0.557 for control vs. hypo) and within bulk (p=0.847 for control vs. K+, p=0.027 for control vs. hypo). **B)** Relative probability of hotspot dynamics (p=0.215 for control vs. K+, p=0.462 for control vs. hypo). **C)** Number of hotspots (p=0.020 for control vs. K+, p=0.189 for control vs. hypo). **D)** Average size of hotspots (p=0.047 for control vs. K+, p=0.794 for control vs. hypo). P-values determined by paired t-test. n=15 paired cells for control and K+, and n=8 paired cells for control and hypo. Grey boxes indicate 95% confidence intervals. Solid black lines indicate mean.

**Fig. 3 – figure supplement 1:**
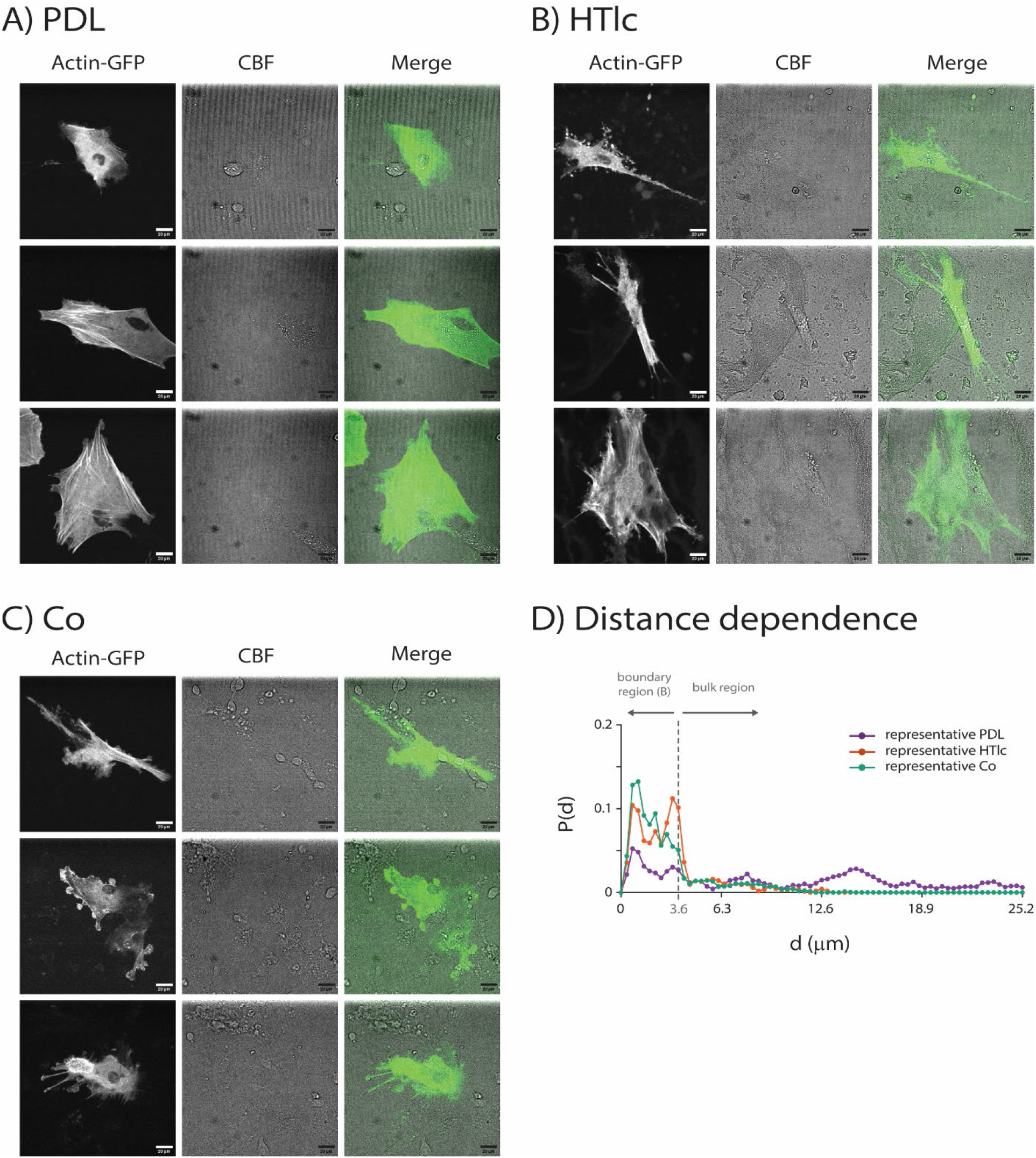
Representative images of primary rat astrocytes in three culturing conditions and supplementary data. **A)** Three representative images of primary astrocytes grown on PDL-coated glass (PDL). **B)** Three representative images of primary astrocytes grown on HTlc nanotopographic films (HTlc). **C)** Three representative images of primary astrocytes co-cultured with neurons (Co) on PDL-coated glass. Left columns show images of astrocytes transduced with actin-GFP from the 488 nm channel. Middle columns show confocal brightfield (CBF) images. Right columns show the merge of 488 nm and CBF channels. Scalebars indicate 20 μm. **D)** Full distance curves for representative PDL, HTlc, and Co cells shown in **Figure 3A**.

**Fig. 4 – figure supplement 1:**
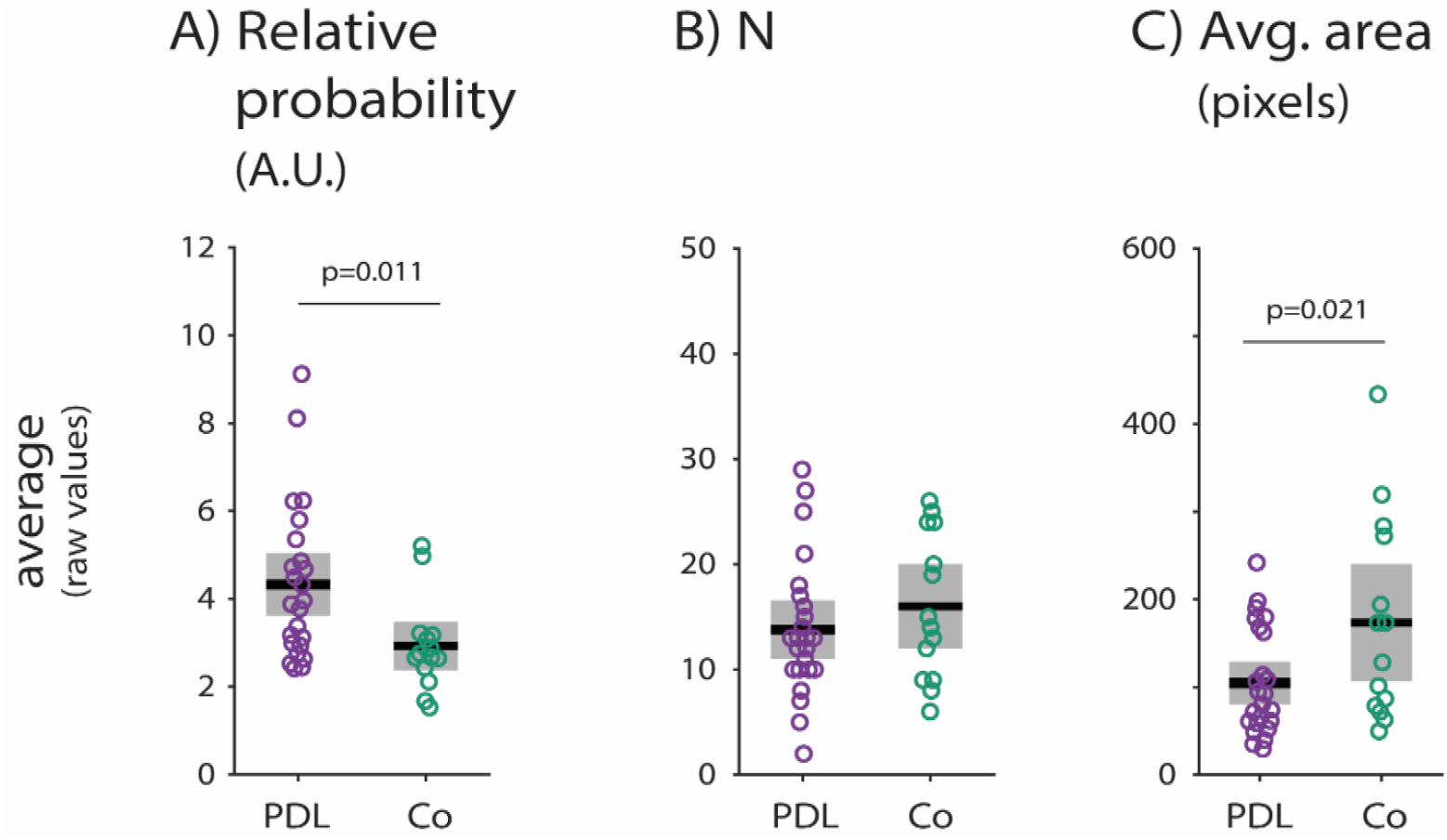
Additional parameters characterizing how the presence of neurons alters actin dynamics within hotspots. **A)** Relative probability of hotspot dynamics (p=0.011 for PDL vs. Co). **B)** Number of hotspots (p=0.333 for PDL vs. Co). **C)** Average size of hotspots (p=0.021 for PDL vs. Co). P-values determined by two-sample t-test. n=24 cells for PDL, and n=14 cells for Co. Grey boxes indicate 95% confidence intervals. Solid black lines indicate mean.

## Notes

### Competing Interest Statement

The authors have declared no competing interest.

### Summary of Updates

Reformatted and with minor changes to Introduction and Discussion.

